# The genetic architecture of local adaptation in a cline

**DOI:** 10.1101/2022.06.30.498280

**Authors:** Fabien Laroche, Thomas Lenormand

**Affiliations:** UMR 1201 Dynafor, Univ Toulouse, INRAE, INPT, EI PURPAN, Castanet-Tolosan, France; CEFE, Univ Montpellier, CNRS, EPHE, IRD, Montpellier, France

## Abstract

Local adaptation is pervasive. It occurs whenever selection favors different phenotypes in different environments, provided that there is genetic variation for the corresponding traits and that the effect of selection is greater than the effect of drift and migration. In many cases, ecologically relevant traits are quantitative and controlled by many genes. It has been repeatedly proposed that the localization of these genes in the genome may not be random, but could be an evolved feature. In particular, the clustering of local adaptation genes may be theoretically expected and has been observed in several situations. Previous theory has focused on two-patch or continent-island models to investigate this phenomenon, reaching the conclusion that such clustering could evolve, but in relatively limited conditions. In particular, it required that migration rate was neither too low nor too large and that the full optimization of trait values could not be eventually achieved by a mutation at a single locus. Here, we investigate this question in a spatially-explicit model, considering two contiguous habitats with distinct trait optima on a circular stepping-stone. We find that clustering of local-adaptation genes is pervasive within clines during both the establishment phase of local adaptation and the subsequent “reconfiguration” phase where different genetic architectures compete with each other. We also show that changing the fitness function relating trait to fitness has a strong impact on the overall evolutionary dynamics and resulting architecture.

## Introduction

Recombination is a key trait for sexual eukaryotes. Genome-wide recombination rates are thought to evolve to limit selective interference and increase the efficacy of natural selection (Otto and Lenormand 2002; Otto 2009). Beyond this global pattern, it has been suggested that low recombination rates within genome could evolve locally to preserve particularly favorable genetic associations. This type of argument applies to sex differences in recombination (Lenormand 2003; Sardell and Kirkpatrick 2020), to inversions capturing loci with strong epistasis (Schwander et al. 2014; Charlesworth 2016), or to loci involved in local adaptation (Pylkov et al. 1998; Lenormand and Otto 2000; Kirkpatrick and Barton 2006). The latter in particular has received a lot of attention, as local adaptation is a ubiquitous phenomenon, and as “genomic islands” of divergence are sometimes, but not always, observed among differentiated populations (reviewed in Nosil et al. 2009; Strasburg et al. 2012). These “genomic islands” may however not result from the evolution of linkage. Instead, they can be relicts of a divergence built by independent evolutionary history of populations. In this case they are simply due to the erosion of this divergence around selected loci (Barton and Hewitt 1989). Because of these different scenarios, the interpretation and evolutionary significance of such genomic islands are difficult to empirically establish. In addition to this empirical difficulty, the theory has not considered the case of spatially explicit models, where dispersal may be distance-limited. Yet, in many situations, local adaptation may take place in a continuous space with a mosaic of different habitats. We investigate this case here.

Local adaptation generally favors lower recombination because migration between habitats leads to positive genetic associations among loci involved in adaptation to these habitats. Lowering recombination rates then preserves combinations of alleles that are locally beneficial, improves the response to selection, and reduces the migration load. Modifier models have been used to make this formal argument (Pylkov et al. 1998; Lenormand and Otto 2000; Lenormand 2012). However, these models do not investigate the specific evolutionary mechanisms by which the linkage between local adaptation loci become tighter. This could occur for several proximal reasons: (1) a local reduction in recombination rate, for instance through the occurrence of a chromosomal rearrangement (Kirkpatrick and Barton 2006; Yeaman 2013), (2) an aggregated recruitment of locally adapted alleles during adaptation to a new environment (Flaxman et al. 2013; Yeaman et al. 2016), (3) or competition between more or less aggregated genetic architectures conferring the same trait values (Yeaman and Whitlock 2011).

Here, we focus on the evolution of aggregated genetic architecture, which encompasses mechanisms (2) and (3). We term mechanism (2) “emergence” and mechanism (3) “reconfiguration”. Aggregation emergence has been studied using either two-patch (Flaxman et al. 2013) or continent-island (Yeaman et al. 2016) models. These models suppose that many loci can contribute to local adaptation to a new environment and they evaluate whether the first mutations contributing to this local adaptation tend to cluster. They conclude that this mechanism is unlikely to strongly bias towards aggregated architecture. This is essentially due to two reasons (Yeaman et al. 2016): first, the linkage to an existing polymorphism affects the establishment probability of a new mutation only within a narrow range of migration rates. Second, this difference in establishment probability is larger for new mutations of small effects that could not establish alone (being swamped by migration) but could establish if tightly linked to a large effect mutation. However, if a distribution of mutation effect size is considered, local adaptation occurs mostly through large effect mutations and this phenomenon therefore plays a minor role. The authors note that this conclusion should also hold in spatially explicit models, such as stepping-stone models, because the region where migration constrains local adaptation is, in addition, geographically limited.

Aggregation reconfiguration has also been studied in a two-patch model (Yeaman and Whitlock 2011), where a trait is under stabilizing selection around different optima in different environments. Many loci are assumed to contribute to this trait, but only a handful of mutations are sufficient to achieve local adaptation in the new environment. Hence, the same phenotypic change can be achieved by very different combinations of mutations at different loci. This allows different genetic architecture to compete, and eventually evolve on the long term. Simulation results show that aggregated architectures evolve in this model but, like for emergence, it occurs in a narrow range of migration rates. There exists a critical migration rate below which a locally favored allele can increase in frequency and contribute to local adaptation. Aggregated architecture evolves roughly between the smallest and largest critical migration rates of plausible mutations, i.e. when migration rates allow polymorphism to persist at local adaptation loci while maintaining substantial gene flow between patches. Like for emergence, aggregation reconfiguration also depends on the distribution of mutation effect size on the phenotype. In particular, if mutations can “stack up” on a single locus, the system can evolve to just one single locus, i.e. a concentrated, rather than aggregated architecture (Yeaman and Whitlock 2011).

Here, we reconsider the evolution of aggregated architecture for three reasons. First, as noted above, it has been suggested that aggregation should be less favored in spatially explicit models with distance-limited migration (at least during the emergence phase). However, this general conclusion is unclear. In particular, the opposite may be true if the condition for local adaptation are less restrictive in spatially explicit models compared to two patch-models. With distance-limited migration, clines form between habitats, irrespective of the relative intensity of migration and selection, provided the geographical scale of habitats is larger than the scale of migration (Nagylaki 1975; Slatkin 1978). Hence, aggregated genetic architecture might evolve within these clines, for a broad range of migration values.

Second, the distribution of mutation effect size seems to play a significant role during the emergence or reconfiguration of aggregation, but several contradictory effects are at play. If mutations of large effect are frequent compared to those of small effect, aggregation is less likely to evolve since fewer loci will be involved in local adaptation and indirect selection will likely play a weaker role in their establishment. At one extreme, if a single mutation that confers perfect adaptation can occur, indirect selection for tight linkage will necessarily play a limited role. Hence, studying aggregation requires imposing some constraint on the maximum phenotypic effect of adaptive loci, hence preventing full ‘stacking’ to occur. Such constraint may also be more representative of the type of genetic variants observed for quantitative traits. Similarly, it would be interesting to clarify the role of small and large effect mutations during the reconfiguration phase. In particular, mutations of large effects could play an ambiguous role in aggregation evolution. They may initially favor aggregation by attracting small effect mutations around them. However, changing genetic architecture around a phenotypic optimum involves crossing a fitness valley, which is expected to be deeper when swapping alleles with larger phenotypic effects. Aggregation during the reconfiguration phase may therefore be slower with mutations of large effects.

Third, the condition for polymorphism at an adaptive locus tightly depends on the shape of fitness trade-off between habitats. In formal one-locus population genetics models, a general case can be considered, where the selection coefficient of alleles in each habitat are free parameters (Nagylaki 1975). Models of polymorphism based on mutations of small effects can consider various fitness trade-off (Ravigne et al. 2009; Débarre and Gandon 2010). Models investigating the evolution of genetic architecture made however more specific assumptions. In continent-island models (Yeaman et al. 2016), no specific trade-off is considered (fitness values are only specified in the island). In multilocus models where the fitness effect of mutations is directly considered (instead of their phenotypic effect), a linear trade-off is often assumed (Flaxman et al. 2013). The fitness effect of alleles simply switch sign across habitats, irrespective of their magnitude. In models using stabilizing selection on a trait, the corresponding shape of the fitness trade-off is implicit, and depends on the specific fitness function considered. This is for instance left as a flexible parameter in Yeaman and Whitlock (2011), but not investigated in details (parameter [] in their eq. 1). With some fitness functions, the ratio of selection coefficient of alleles across habitats depend on the magnitude of the phenotypic effect of the mutation and on the current mean trait value of the population (compare Gauss and Laplace fitness function sketched on figs. 1 and 2). Hence, changing the fitness function may affect the relative contribution to local adaptation of mutations of large or small phenotypic effects, as they will exhibit different fitness asymmetries across habitats. Furthermore, these relative contributions may also change dynamically during the process of adaptation, especially when those fitness asymmetries depend on the current mean trait values in the population. Hence, the choice of the fitness function will likely influence the initial and long-term dynamics of aggregation.

**Figure 1.**
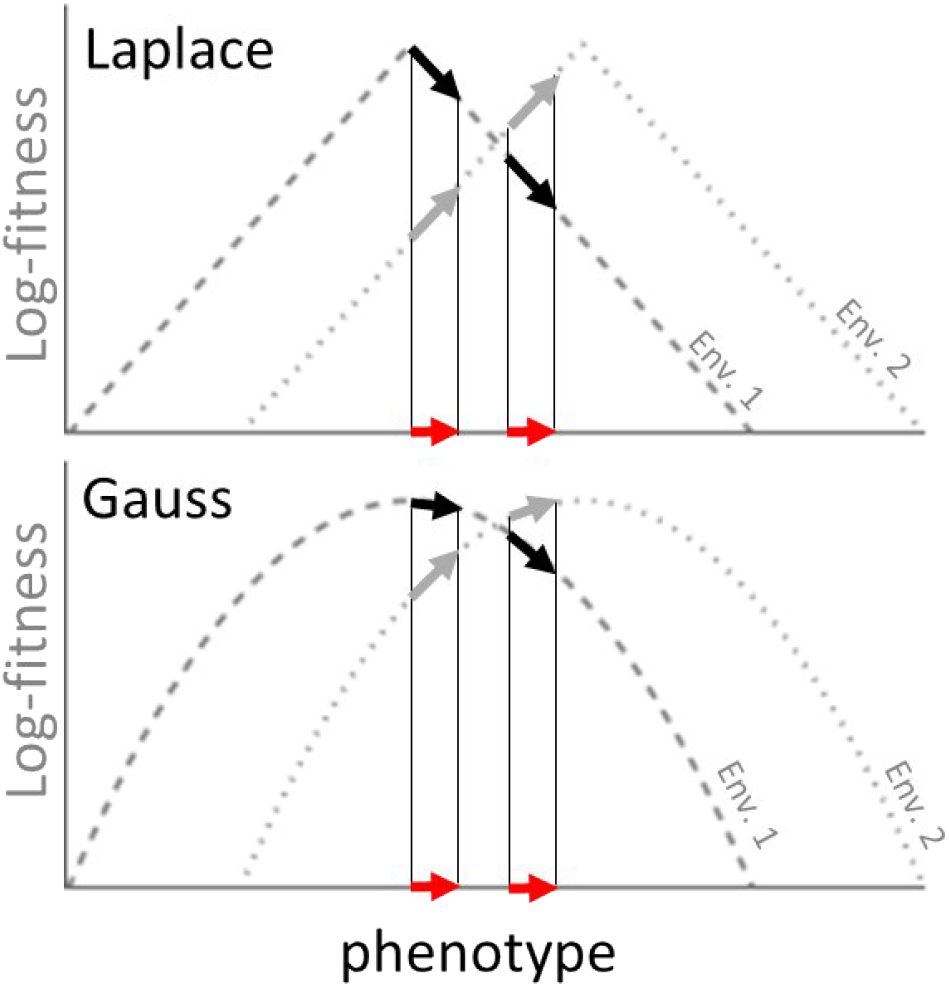
Laplace and Gauss fitness functions. The red arrow represents a mutation increasing the trait value. With Laplace fitness function, the log-fitness gain of the mutation in environment 2 (gray arrow) is equal to the log-fitness loss in environment 1 (black arrow), irrespectively of the current phenotype (in absence of optimum overshoot). With Gauss fitness function, the log-fitness gains (and losses) depend on current phenotype.

**Figure 2.**
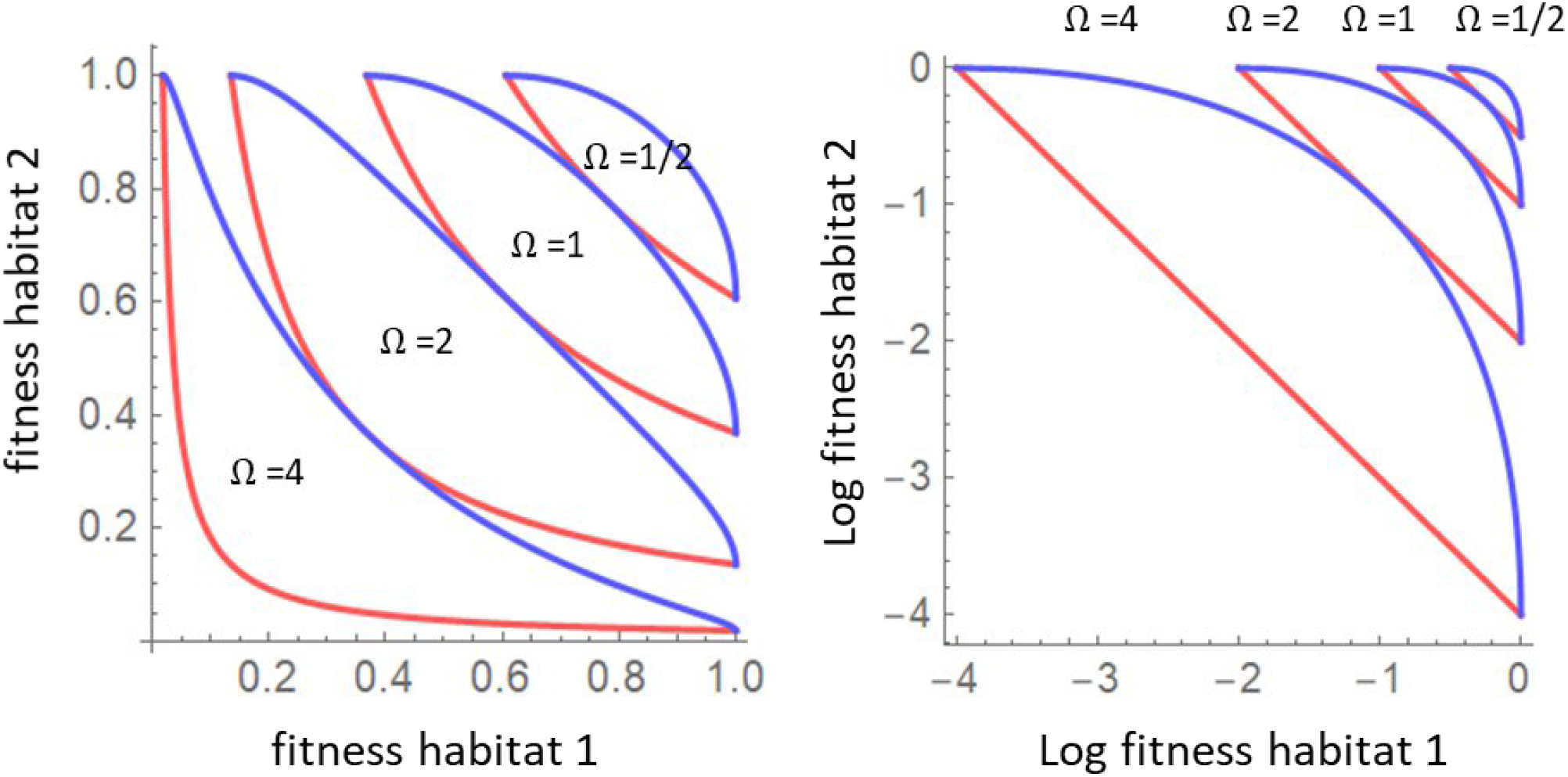
Fitness trade-off curves between environments, for different intensity of stabilizing selection (*Ω*). Without loss of generality, the difference in optimal phenotype between habitats is set to 1. Red: Laplace fitness function ([] = 1). Blue: Gauss fitness function ([] = 2). Left : fitness; Right: log fitness.

To address these issues, we investigate how genetic architecture evolves in a spatially continuous model with a small “pocket of adaptation”. This spatial setting stands as the simplest extension of the continent-island or two-patch models of adaptation considered in previous studies cited above to a spatially-explicit context. We consider two habitats in a one-dimensional circular stepping stone. We consider stabilizing selection on a trait, and we consider that only two kinds of mutations can occur, with either a small or a large phenotypic effect. These effects are fixed and therefore a full stacking of effects at a single locus is not possible. This allows us to clearly isolate the role of mutations with different effect sizes on the evolution of aggregated architectures. Last, we consider two types of fitness functions (Gauss and Laplace) to investigate whether the fitness function and the underlying shape of fitness trade-off matter for the evolution of aggregated architectures.

## Methods

### Model

We model the process of adaptation to a “pocket” (i.e. a finite, contiguous area) of habitat. We use a model with discrete time and space. We consider a one-dimensional landscape made of *n* patches regularly positioned on a circle, each with a constant number of individuals *N*. Without loss of generality, we assume that the distance between two consecutive patches is one distance unit. This landscape is made of two habitats : *n*-*l* contiguous patches correspond to habitat 1, and *l* patches to habitat 2. Individuals express a phenotype *z*, which is under stabilizing selection around a different optimum value *z*_*opt*_ in the two habitats. We denote *z*_*opt*_(*i*) the optimum in patch *i*. We assume that *z*_*opt*_(*i*) = 0 if patch *i* is in habitat 1 and *z*_*opt*_(*i*) = 1 if patch *i* is in habitat 2 (fig. 3A). We suppose that an individual fitness in patch *i*, denoted *w*_*i*_(*z*), decreases with the deviation of the trait *z* from *z*_*opt*_(*i*) :

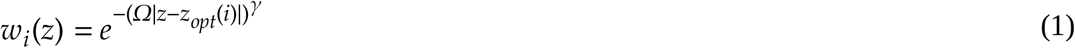

where *Ω* is the intensity of stabilizing selection (which we assume constant across all patches), and *γ* controls the shape of the fitness function (following Yeaman and Whitlock 2011). We will focus on two cases: *γ* = 2, which is the widely used Gauss function and *γ* = 1, which we will refer to as the Laplace function (fig. 1). An important difference between these two cases is that a phenotypic change *ε* around *z*_*opt*_ has a negligible effect on fitness at first order in the Gauss case (where *w*(*z*)=1 + *O*(*ε*^2^)), but not in the Laplace case (where *w*(*z*)=1 - *Ω*|*ε*| + *O*(*ε*^2^)). These fitness functions also implicitly determine the shape of fitness trade-off across environments (fig. 2).

**Figure 3.**
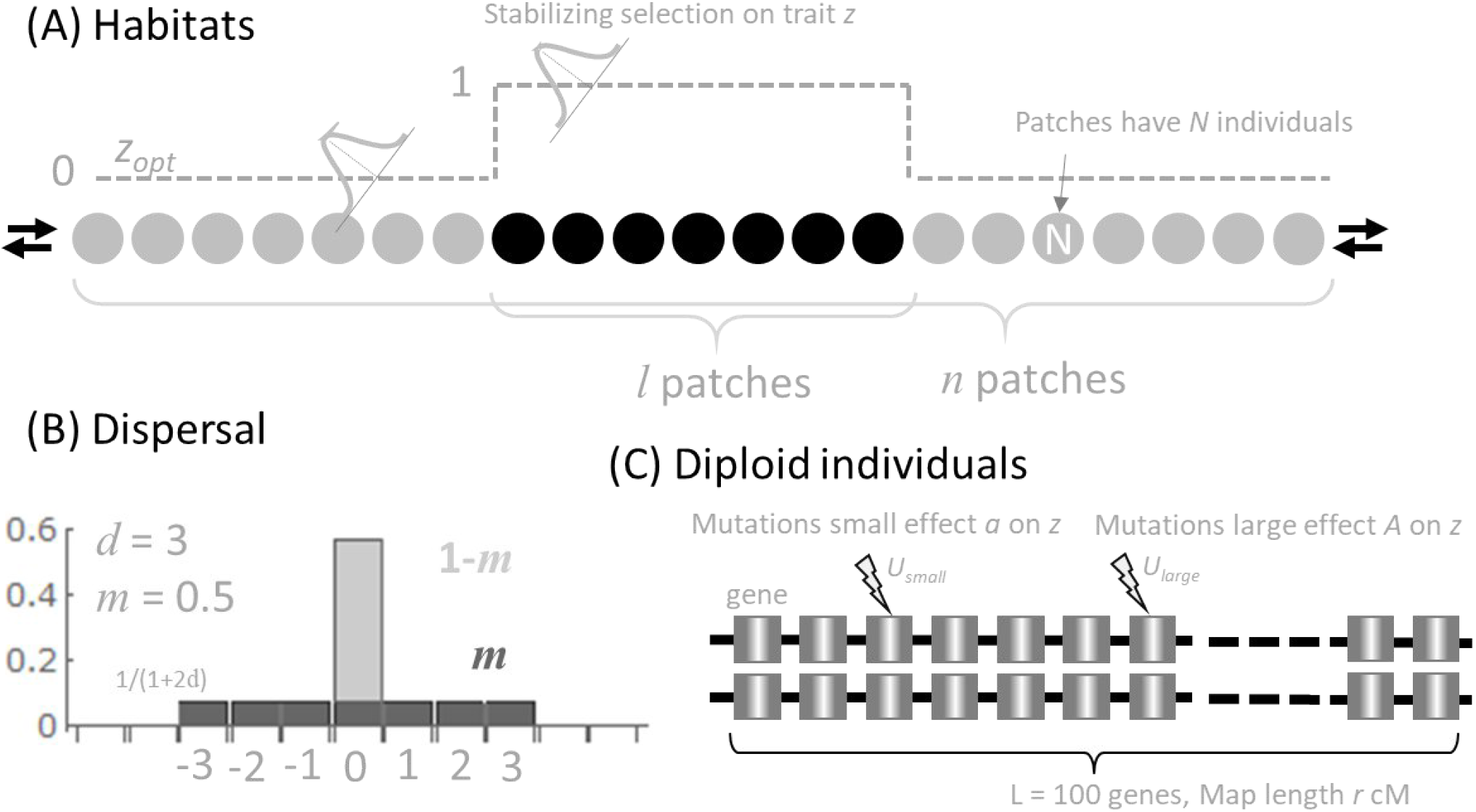
Model assumptions. (A) The landscape is one-dimensional and circular with *n* patches with *N* individuals. Habitat 1 and 2 are made of *n*-*l* and *l* contiguous patches with an optimal trait value *z*_*opt*_ = 0 and 1, respectively. Trait *z* is under stabilizing selection around this optimum. (B) Dispersion kernel. A proportion 1-*m* of individuals stay in their patch, and a proportion *m* is uniformly distributed among patches within distance *d* of the focal patch (focal patch included). Hence, each non-focal patch receives *m*/(1+2*d*) migrants. (C) Individuals are diploid and have a pair of chromosomes with *L* loci determining trait *z*. Mutation of small or large effects can occur on all these loci.

We modeled diploid individuals carrying two chromosomes with *L* evenly spaced loci determining the trait *z* (fig. 3C). The life-cycle of the model begins at juvenile diploid stage. The number of juveniles is assumed very large, non-limiting in the dynamics, and equal among patches. Migration of juveniles occurs according to the kernel *Π* illustrated on fig. 3B. *Π* is a *n*×*n* matrix, and *Π*_*ij*_ is the proportion of juveniles from patch *j* that reaches patch *i* after migration. This kernel depends on two parameters: the proportion of migrants leaving a patch (*m*) and the maximal migration distance (*d*). The variance of this kernel is *mσ*^*2*^ where *σ*^*2*^ = *d*(*d*+1)/3.

Juveniles are then regulated to *N* mature individuals in their arrival patch (i.e. *N* mature individuals are randomly sampled within the local juvenile pool, the other juveniles die). Mature individuals produce a large, non-limiting number of gametes through meiosis. Selection acts on the relative chance that an adult reproduces, which is proportional to its fitness *w*_*i*_(*z*). The number of cross-over at meiosis follows a Poisson distribution of parameter *r*, the map length. Cross-over events are then uniformly distributed on the chromosome. Mutation can also occur on loci at meiosis. Each locus is tri-allelic, with effect 0, *a* or *A* on *z*, with 0 < *a* < *A*. Effects of alleles and loci are additive on *z*. Small effect mutations (of effect *a*) reversibly occur at rate *μ*_*a*_ (mutation events 0→*a, a*→0 occur with probability *μ*_*a*_). Large effect mutations (of effect *A*) reversibly occur at rate *μ*_*A*_. Mutation events *a*→*A, A*→*a* do not occur, and we consider that large effect mutations occur less frequently than small effect mutations (typically *μ*_*A*_ = *μ*_*a*_/1000; see Table 1).

**Table 1.** Parameter values used in simulations. List of parameters values used in the simulations. We used fourteen values of *l*, for both long term and initial phase of adaptation, for both Laplace and Gauss fitness functions. We used 20 replicates per combination of parameter values. This resulted in 14 × 2 × 2 × 20 = 1120 simulations. For each simulation with *l* ≥ 21, we performed a control simulation where habitat 1 was removed, which resulted in 9 × 2 × 2 × 20 = 720 control simulations. These control simulations were not performed for *l* < 21, because dispersal kernel with *d* = 10 (the value used throughout) was undefined in that case.

| Parameter | Definition | Values |
| --- | --- | --- |
| $N$ | Total number of patches | 400 |
| $L$ | Number of patches of habitat 2 | 1,5,9,...,21, 29, 37,..., 85 |
| $N$ | Number of individuals within patches | 100 |
| $M$ | Proportion of migrating juveniles | 0.5 |
| $d$ | Maximum dispersal distance of migrants | 10 |
| $\gamma$ | Shape of fitness around local optimum | 1 (Laplace), 2 (Gauss) |
| $\Omega$ | Intensity of selection | 1 |
| $L$ | Number of loci on the chromosome | 100 |
| $a$ | Effect size of small mutations | 0.01 |
| $A$ | Effect size of large mutations | 0.1 |
| $\mu_a, \mu_A$ | Mutation rates towards alleles of small and large effects | 1e-5, 1e-8 |
| $r$ | Expected number of recombination events at meiosis | 0.5 |
| $T$ | Number of generations in a simulation | <50'000 (initial phase), 400'000 (long term) |

Mature individuals are hermaphroditic and randomly mate within their patch. Diploid juveniles of the next generation in a patch are thus obtained by randomly sampling two gametes in the patch gamete pool. As mentioned earlier, the total number of juveniles produced is large, non-limiting and assumed equal among patches, which corresponds to a soft-selection regime (selection acts within patches and does not change the number of adults per patch; Karlin 1976).

We consider a population initially fixed for the wild type alleles 0, and we follow adaptation to a new pocket of habitat 2 which requires the occurrence and spread of these small or large effect alleles *a* or *A*. We consider the case where there are more available loci than required to reach the new optimum *z*_*opt*_ = 1 in habitat 2. For instance with *a* = 0.01, *A* = 0.1, the optimal phenotype can be reached by either 5 loci fixed for large effect alleles or 50 loci fixed for small effect alleles (or any intermediate combination). If *L* = 100, as we typically use, there are thus more available loci than required to reach the optimum in habitat 2, irrespective of the combination of large and small effect alleles used. We can therefore study whether the genetic architecture evolves to be “aggregated”, i.e. whether loci contributing to the phenotypic change in habitat 2 are closer from each other on the chromosome than expected by chance.

### Simulations

For simulations, we use a Wright-Fisher approach, drawing each new individual from the previous generation. First, we determine the patch of the parents (according to the migration kernel). Each parent is then sampled with replacement from this patch, weighted by its fitness. One gamete is sampled from each parent (with possible recombination and mutation events) to form the new individual and this is repeated to reach *N* individuals in each patch. The corresponding code is written in Java programming language and source code is available at 10.5281/zenodo.6602165.

Table 1 gives the parameter values used in the simulations. We consider a landscape made of *n*=400 patches, which allows browsing a wide range of proportions of habitat 2 from 0.25% (i.e. *l*=1) to 21% (i.e. *l*=85). We consider *N*=100 individuals per patch, which is a compromise between minimizing the effect of drift and keeping computational time moderate, and is also a reasonable population size for many organisms. The intensity of stabilizing selection *Ω* is set equal to 1, meaning that when an individual perfectly adapted to one habitat is moved to the other habitat, it has a relative fitness of e^-1^=0.36 compared to locally adapted individuals (irrespective of the considered shape of fitness function). The number of loci is set to *L=*100, here again making a compromise between the need of leaving room for variation in the genetic map to occur and the need to keep computational time moderate. The values used for the proportion of migrants leaving a patch *m*=0.5 and the maximal migration distance *d=*10 correspond to organisms with quite high propensity to leave the home patch but distance-limited dispersal. This can for instance occur when patches correspond to a micro-habitat where dispersing is mostly determined by the propensity to disperse, with little influence of internal movement within the patch (see Jonsson 2003 for an example of high propensity to disperse in a fungicolous beetle). Considering *d* = 10 ensures that, for the lower bound of habitat 2 sizes, many individuals from habitat 1 can immigrate at the center of habitat 2, while for the upper bound, it does not occur.

Simulations are initialized with all individuals having phenotype *z* = 0. We focus on two timescales, one corresponding to the initial phase of adaptation (to follow the emergence of aggregation), the other considering adaptation in the long term (to follow the reconfiguration phase). The initial phase of adaptation is defined as the period between initialization and the first time when the average phenotype in the patch located at the middle of habitat 2 reaches 90% of its long term average stationary value. This initial phase lasted typically less than 50’000 generations in our simulations. The long term phase extends beyond this initial phase, up to 400’000 generations. During this phase, we always eventually visually observe a stationary regime in average phenotype in the patch located in the middle of habitat 2. The length of our simulations is sufficiently large to ensure that this stationary regime represents at least 75% of the simulation time. These definitions of initial and long-term phases are somehow arbitrary, and we cannot discard the fact that waiting longer in the long-term phase could reinforce the aggregation patterns reported later in the text. However, these choices are sufficient to observe marked differences between the emergence and reconfiguration phases, which is our objective. The frequency at which the state of the system is recorded differs between the emergence (one record every 500 generations) and the reconfiguration phases (one record every 4000 generations).

### Measure of aggregation

In a given patch *i*, denote *v*_*i*_ *=* (*v*_*i*1_,*v*_*i*2_,…,*v*_*i*L_) the profile of the frequencies of alleles with effect size *a* (=0.01 in our study) along the *L* loci. If 50 loci are fixed for small alleles, and that all these contributing loci are next to each other, *v*_*i*_ has a “square” profile: equal to 1 on a range of 50 contiguous loci and to 0 everywhere else (with eight loci it would look like “11110000”). By contrast, if the 50 loci are randomly scattered along the chromosome, vector *v*^*i*^ looks like a “white noise” along the *L* loci (random permutations of 0 and 1).

Many possible metrics can be used to measure aggregation of alleles frequencies along the profile *v*_*i*_. We opt for a measure based on a spectral analysis, making an analogy between the profile *v*_*i*_ along the chromosome and a time series with *L* time steps. The spectral density of *v*_*i*_ quantifies the contributions of various “frequencies along the chromosome” (FAC below) to the profile. Coming back to the two extreme examples of profile given above, a square wave has a spectral density with a marked peak for low frequencies. In other words, low FACs contribute a lot to the profile, and therefore their weighed average is lower than if all FACS were equally contributing. By contrast, a white noise has a flat spectral density across all the spectrum. Hence, we propose that aggregation can be measured by the bias of the spectral density towards low FAC values relative to white noise baseline.

To build a metric corresponding to this idea, we first compute the spectral density *s*_*f*_ of the FAC *f* using a discrete Fourier transform of *v*_*i*_. We then measure a weighted mean of FAC according to their contribution to the profile:

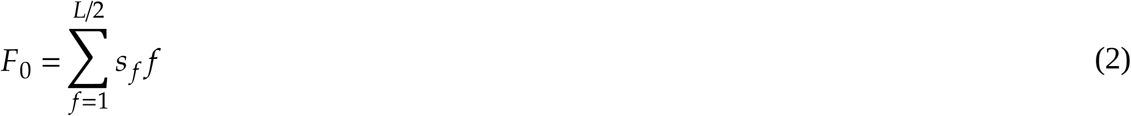

Low *F*_0_ indicates that the spectral density is biased toward low signal frequencies. We scale *F*_0_ between the maximal (*F*_*max*_) and minimal (*F*_*min*_) values possible to achieve for a vector with the same elements than *v*_*i*_, but in a different order. *F*_*max*_ measures white noise and is computed as an average over random permutation of *v*_*i*_. *F*_*min*_ measures a proxy of the maximal aggregation that could be reached, and is computed on a vector where all values in *v*_*i*_ are sorted. The scaled measure of aggregation is then obtained as:

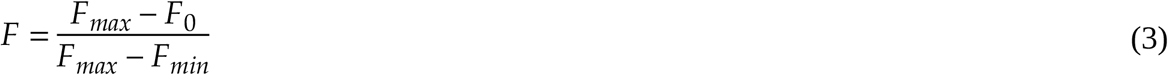

It is close to 0 when there is no aggregation (the weighed signal frequency is close to the value obtained on random permutations), whereas it is close to one when aggregation is maximal (close to a square signal). We generate permutations of frequencies’ position in *v*_*i*_ and compute a p-value to assess whether the observed value of the metric is significantly higher than expected under the hypothesis of random positioning.

This aggregation metric can be obtained using the frequency vector of small effect alleles, but it can be similarly applied to the frequency vector of large effect alleles alone, or to vector of the sum of small and large alleles frequencies.

## Results

### Condition for polymorphism

We first consider the deterministic condition for the existence of a cline for an allele introduced in a monomorphic population where individuals have trait value *z*. We consider a mutation at one locus creating a new phenotype *z*_*mut*_ (i.e. the mutation effect size is *z*_*mut*_ *-z*). Assuming that the adult population size *N* is large within each patch, the gamete frequency of a rare mutant within the *n* patches *x*(*t*)=(*x*_1_(*t*), *x*_2_(*t*), …, *x*_*n*_(*t*)) follows the linear recursion (Karlin, 1976):

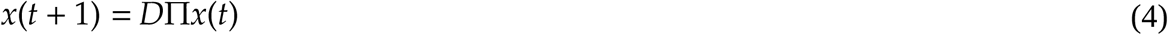

where *D* is a *n*×*n* diagonal matrix such that *D*_*ii*_ = *w*_*i*_(*z*_*mut*_)/*w*_*i*_(*z*). The mutant can invade the resident population if and only if the dominant eigenvalue of *DΠ* is above 1.

Denote *λ*(*x,y*) the dominant eigenvalue of *DΠ* when *z* = *x* and *z*_mut_ = *y*. Linearizing for small phenotypic effects of mutants (i.e. with *z*_*mut*_ *-z = ε* with *ε →* 0), then, for *0 < z < 1*, we have:

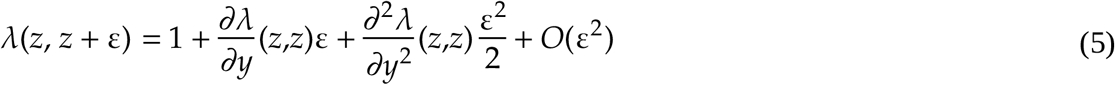

In appendix A, we give a general expression of this dominant eigenvalue in terms of the parameters of the model. For simplicity, here, we only focus on the ‘semi-infinite’ limit, where the size of habitat 2 is kept constant, while the size of habitat 1 is considered infinitely large (*n*→+∞, Nagylaki 1975). Defining *s*_*1*_ (resp. *s*_*2*_) the value of the selection gradient *w*_*i*_*’*(*z*)/*w*_*i*_(*z*) in patches of habitat 1 (resp. 2), and *t*_1_ (resp. *t*_2_) the value of *w*_*i*_*’’*(*z*)/*w*_*i*_(*z*) in patches of habitat 1 (resp. 2), we obtain:

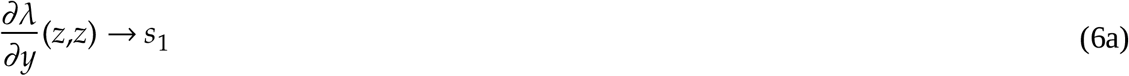

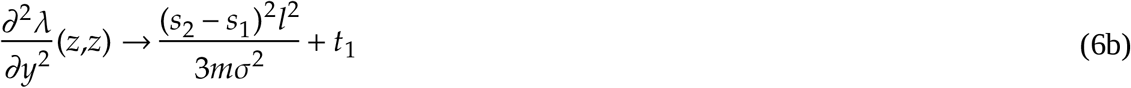

Using equations (6a,b) in equation (5) yields the general condition for the existence of a cline in this model. Results for the Laplace and Gauss fitness functions are presented below and summarized in fig. 4 (details of calculus are provided in Appendix B).

**Figure 4.**
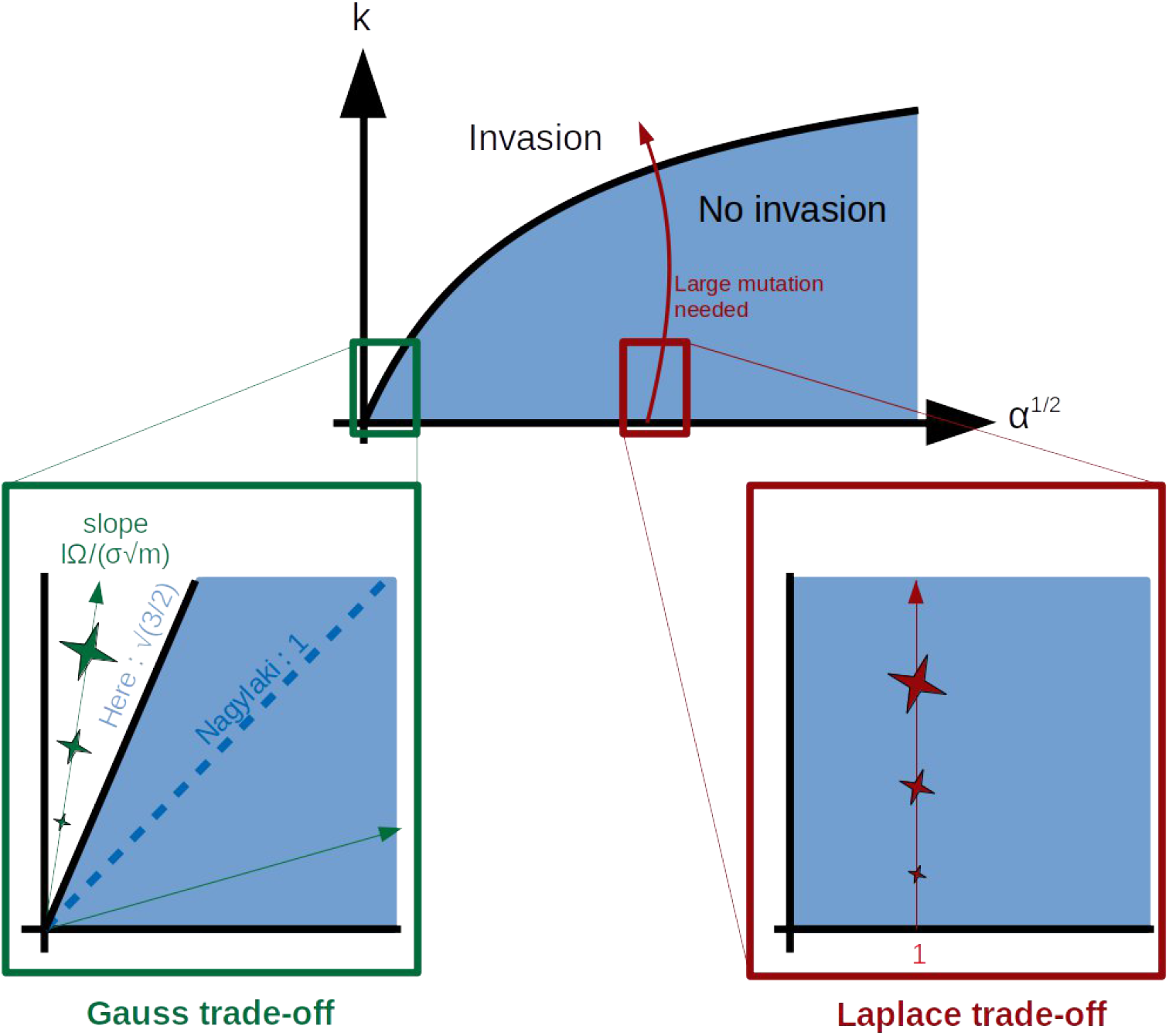
Conditions for polymorphism in the semi-infinite limit of our model. The upper panel correspond to the condition for the existence of a cline in an habitat ‘pocket’ as predicted using (Nagylaki 1975). See Lenormand (2002), Box 2, fig. II. The figure shows whether a mutant can invade (white area) or not (blue area) the *z*\*=0 resident phenotype as a function of two compound parameters *k*, and *α* (if it does, a cline establishes between the habitats). On the y-axis *k* measures the relative scales of the spatial heterogeneity (width of the pocket) and of the ‘characteristic length’ (σ/√s), which weighs the strength of selection relative to gene flow. On the x-axis, *α*^1/2^ measures the square root of the ratio of selection coefficients outside/inside the pocket. The lower panels zoom on areas of the figure where small mutants are positioned with a Gauss (green, left) and Laplace (red, right) fitness trade-off. In both panels, mutation with increasing phenotypic effects are presented using stars with increasing size. In the Gauss case, mutations of different effect sizes follow lines with zero intercept and slopes that depend on the parameters (see eqs. 11, 12a, 12b). Mutants can invade if this slope is above the critical slope corresponding to the frontier between the blue and white areas. This is an all-or-nothing situation. Mutations of small and large effects can either all invade or none of them can invade. In the Laplace case, mutations are on a vertical line with *α* = 1 (symmetry between habitats). Infinitesimal mutations cannot invade, but large mutations can form clines if their phenotypic effect is sufficiently large.

We use Nagylaki’s 1975 results as a comparison. In Nagylaki’s model, space is continuous and migration is modeled as a diffusion process. The key result is that the condition for the existence of a cline is determined by two combined dimensionless parameters (see box 2, fig. II in Lenormand 2002; fig. 4 here). The first parameter, noted *k*, measures the scale of the spatial heterogeneity (here the size of the pocket, *l* in eq. 7a) relative to the ‘characteristic length’ (Slatkin 1973) of migration scaled by selection intensity. By selection intensity, we mean the selection coefficient of target mutant with phenotypic effect *ε*, denoted *S*_1_(*ε*) (resp. *S*_*2*_(*ε*)) in habitat 1 (resp. 2). In practice, *S*_*i*_(*ε*) = *w*_*i*_(*z+ε*)/*w*_*i*_(*z*) - 1 for i=1,2. Specifically, the characteristic length *k* is the ratio of the standard deviation of parent offspring distance (from the dispersal kernel, 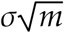 in our case) divided by the square root of the selection intensity in habitat 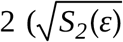 in our case), which yields 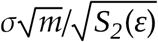 in eq. 7a). The second parameter, noted *α*, measures the fitness asymmetry between habitats. In the context of our study, the two dimensionless parameters defined in Nagylaki (1975) are expressed as follows:

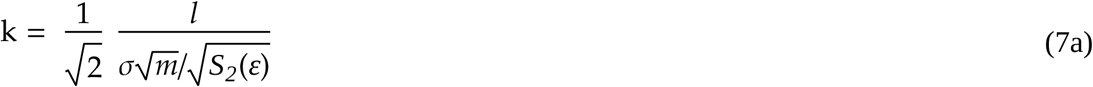

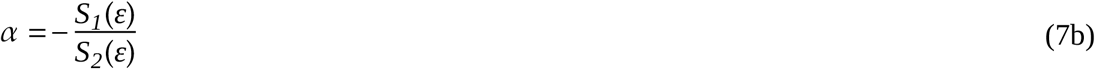

Here we focus on the case where *z*=0 (resident is fully adapted to habitat 1) and *ε* > 0. The fitness asymmetry between habitats *α* is thus positive : the mutant allele has a fitness trade-off among habitats. The existence of a cline is not relevant in other cases.

#### Laplace trade-off

When *γ* = 1, (6a) simplifies to *∂λ/∂y*(*z,z*) = - *Ω* irrespective of *z* value in] 0,1[. Because *∂λ/∂y*(0,0_*+*_) = - *Ω, z* = 0 cannot be invaded by infinitesimal mutations. In this case, Nagylaki’s dimensionless parameters of equation (7a,b) are:

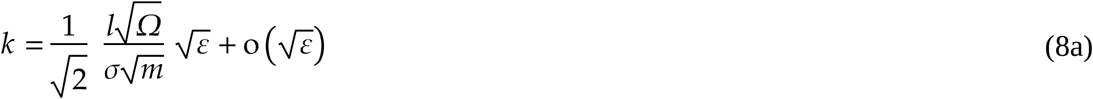

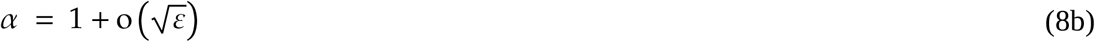

Equations (8a,b) allow locating the Laplace trade-off on Nagylaki’s chart (fig. 4) : infinitesimal mutations cannot invade unless some limit case is considered with respect to other parameters. We consider the limit case where the standard deviation of the dispersal kernel is small compared to the size of habitat 2 (i.e. 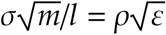 with *ρ* a constant) and we assume that taking this limit does not generate new terms of order *ε* or larger in equation (5). Then, using equations (6a) and (6b), equation (5) becomes:

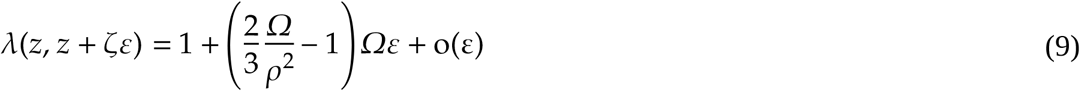

and polymorphism can arise from *z\**=0 if 2*Ω*/(3*ρ*^2^) > 1 which can be reformulated as a minimum requirement (a “swamping limit”) for the size of habitat 2:

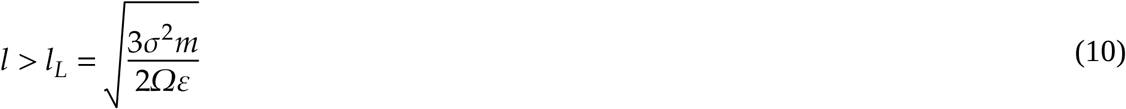

Note that the swamping limit *l*_*L*_ decreases as the effect size of the mutation (*ε*) increases (in other words, large effect mutations can invade in a broader range of parameter values, including smaller pockets of habitat 2). We therefore use the notation *l*_*L*_(*ε*) for the swamping limit of a mutation with effect size *ϵ* in the Laplace case.

#### Gauss trade-off

When *γ* = 2, (6a) simplifies to *∂λ/∂y*(*z,z*) = - 2*Ω*^2^*z*. Therefore, contrary to the previous case, *∂λ/∂y*(0,0_*+*_) = 0, which entails that *z*=0 can be invaded by small mutations if *∂*_*2*_*λ/∂y*_*2*_(0,0_*+*_) > 0. Using equation (6b), this condition becomes:

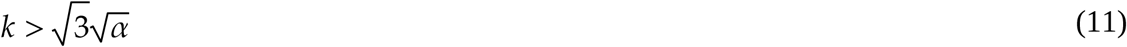

where Nagylaki’s parameters are here equal to:

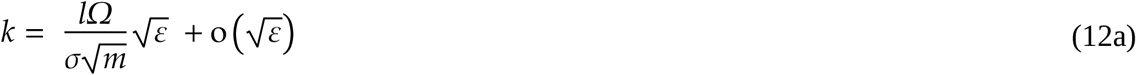

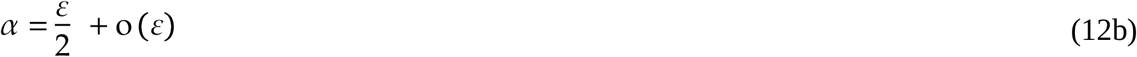

This result (eq. 11) is close to Nagylaki’s (1975; eq. 38 and 40a) one, where, for small *α*, the condition for polymorphism is 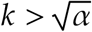. An important difference with the Laplace case is that the asymmetry between habitats depends on the mutation effect size (eq. 12b). Mutations of smaller phenotypic effects are exposed to weaker selection but they enjoy a more favorable outer/inner ratio of selection coefficients compared to mutations of larger phenotypic effects. At the limit, for mutations of very small phenotypic effect, this ratio tends towards zero (eq. 12b), meaning that mutations of very small effect enjoy an advantage in habitat 2, while suffering from virtually zero fitness decrease in habitat 1. This is caused by the fact that the Gauss fitness function is flat around the optimum. The consequence is that either all mutations can invade or none, depending on whether 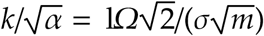) is greater or lower than 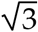, respectively. In other words, a polymorphism arises when the size of habitat 2 is above the swamping limit *l*_*G*_ :

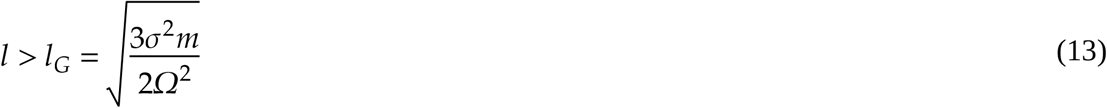

but this does not depend on the phenotypic effect sizes of mutations.

### Contribution of small and large effect mutations to local adaptation

Comparing the swamping limit in the Gauss case (eq. 13) with our approximation of swamping limits in the Laplace case (eq. 10) for the effect sizes considered in our study (*a*=0.01, *A*=0.1; Table 1), we expect that: *l*_*G*_ < *l*_*L*_(*A*)<*l*_*L*_(*a*). When *l* < *l*_*G*_, no polymorphism should emerge, irrespective of the shape of the fitness trade-off.

When *l*_*G*_ < *l* < *l*_*L*_(*A*), we expect a polymorphism to emerge with a Gauss fitness trade-off, but not with a Laplace trade-off. In this case, the condition for maintaining polymorphism does not depend on the effect size of mutations. Hence, as far as mutations of small effect *a* occur more frequently than mutations of large effect *A*, they should strongly contribute to the phenotypic change. However, the stochastic loss of beneficial mutations that occur when mutations are present in few copies should nevertheless penalize mutations of small effect. To quantify these antagonistic effects, we can compare the rate of establishment of small versus large mutations in the Gauss case. Using Haldane (1927) approximation, a mutant allele of phenotypic effect *ε* has a probability of establishment that is approximately 2(*λ*(*z*,z*+ ε*) - 1). Therefore, after a period of *t* generations, when neglecting interactions among alleles and the global shift of background phenotype (i.e. all the alleles are still rare), we expect the number of established small and large effect alleles to be proportional to μ_a_(*λ*(*z*,z*+a*) - 1)*t* and μ_A_(*λ*(*z*,z*+A*) - 1)*t*, respectively. Among established alleles, the proportion *P*_*A*_ of large effect mutations should then be approximately:

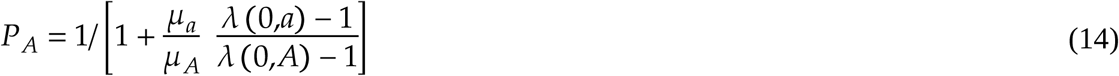

and the proportion *Q*_*A*_ of trait evolution z(t)-z(0) within habitat 2 due to alleles with effect size *A* should be approximately

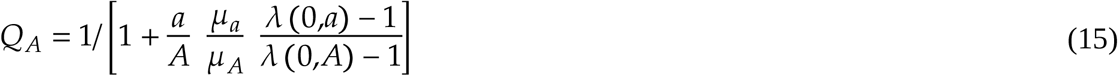

In the Gauss case,

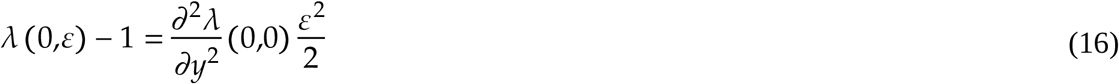

Hence, when *l>l*_*G*_, the proportion of trait evolution z(t)-z(0) within habitat 2 due to alleles with effect size *A* should be approximately

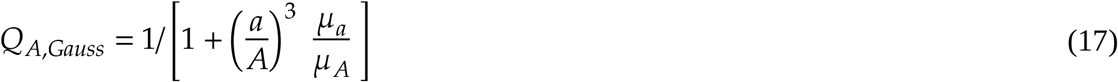

which is constant and equal to 0.5 with the parameter values used in our simulations (Table 1). Comparing Gauss and Laplace cases we thus obtain that, when *l*_*G*_ < *l* < *l*_*L*_(*A*), a polymorphism arises in the Gauss but not in the Laplace case. In the Gauss case, the relative contributions of small and large effect mutations is given by equation (17). When *l*_*L*_ *A* < *l* < *l*_*L*_ *a*, the situations remain unchanged in the Gauss case, but a polymorphism arises in the Laplace case, only involving large effect alleles. When *l*_*L*_ *a* < *l*, the situations remain unchanged in the Gauss case, and a polymorphism arises in the Laplace case, involving both large and small effect alleles. In the latter, using equations (9, 10) and the expression of ρ in equation (13), the proportion *Q*_*A,Lap*_ of trait evolution z(t)-z(0) within habitat 2 due to alleles with effect size *A* is:

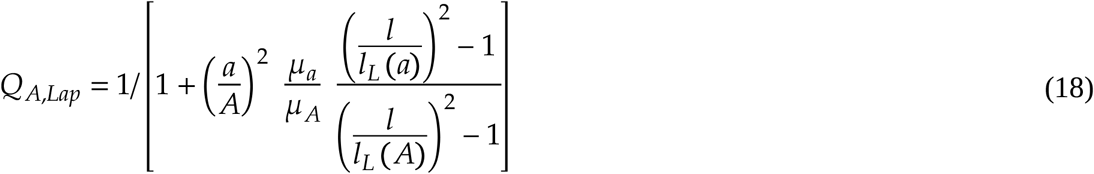

Equation (18) shows that as the size of habitat 2 increases from *l*_*L*_ *a, Q*_*A,Lap*_ decreases from 1 (only large alleles A contribute to the phenotype) to *Q*_*A,Gauss*_ (the contribution found in the Gauss case).

In summary, the emergence of a polymorphism requires a larger size of habitat 2 in the Laplace (*l*_*L*_(*A*)) than in the Gauss case (*l*_*G*_). When the size of habitat 2 is above the critical size *l*_*L*_(*A*), large effect alleles A should make a stronger contribution to the phenotype in the Laplace compared to the Gauss case during the emergence phase, and this difference should decrease towards 0 as habitat 2 becomes large.

#### Effect of the “background trait value” -

These relative contributions of mutations of small and large effects are computed initially, when the average phenotype in habitat 2 just begins to evolve towards its optimal value. However, during this process, a cline builds up, and the average trait value increases in habitat 2, but also in habitat 1, at least close to the boundary between habitats. In the Laplace case, this is not strongly consequential, as the fitness function is linear and its local slope does not depend on average trait values, at least until adding one mutation causes a phenotypic overshoot in habitat 2. However, this is more consequential in the Gauss case: the slope of the Gauss fitness function in habitat 1 gets steeper as trait value increases (instead of being flat initially, see fig. 1). When the average trait value increases in the Gauss case, mutations of small effect no longer enjoy the benefit of having more favorable outer/inner ratio of selection coefficients. As we explore in appendix B, this favors the establishment of mutations of larger effects compared to the initial expectation, during the emergence phase. In particular, when the average trait value becomes larger than a threshold value, small alleles cannot invade anymore and only large alleles can pursue the adaptative process (until the phenotype becomes close to the optimum so that overshooting becomes a problem). Overall, this effect of “background trait value” suggests that equation (17) may underestimate the contribution of large effect mutations to adaptation in the Gauss case. This bias should become lower when the size of habitat 2 increases, as counter-selection outside habitat 2 becomes less and less important in this case.

#### Effect of “aggregation” -

Last, the establishment rates computed above do not account for indirect selection and linkage disequilibria among several alleles. In the Laplace case, for a given pocket size, only mutations exceeding a given effect size can form a cline and contribute to local adaptation. However, we also expect that during the establishment of local adaptation, some small effect mutations that occur in close linkage (to one another or to large effect mutations) could also invade. At small timescales, this effect will mostly concern small effect mutations, that co-occur much more frequently than large effect mutations (assuming, as we do, that *µ*_*a*_ ≫ *µ*_*A*_). For instance, for a pair of mutations, cases of co-occurrence will scale with the square of mutation rate, which can be limiting at short time scales (Yeaman 2013). This aggregation bias should be particularly strong below the swamping limit of small effect mutations, where mutations of small effect can establish only if they are aggregated. This effect will also favor small effect mutations in the Gauss case by increasing the establishment probability of co-occurring small-effect mutations that happen to be in close linkage as background trait value increases. Overall, this effect of “aggregation” favors small effect mutations in both the Laplace and Gauss cases. It should lead to cluster of small effect mutations, possibly grouped around a large-effect mutations. Here too, this bias should decrease with increasing size of habitat 2, as indirect selection only matters within clines at the boundary between habitats, i.e. in a fraction of the landscape that becomes relatively less important when the size of habitat 2 increases.

### Simulations of the short-term emergence phase

#### Laplace trade-off

Our analytical computation of the invasion threshold for large alleles derived from equation (10) is remarkably accurate (fig. 5A): below this critical threshold, there is no adaptation in our simulations. By contrast, we observe that small-effect mutations contribute significantly to the establishment of local adaptation even below their predicted swamping limit. As expected, they show a clear signal of aggregation in this case, and this signal is stronger when computed on small and large effect mutations combined (fig 5B). In fact, small-effect mutations form clusters grouped around large effect mutations that establish first. Fig. 7A shows a typical example of this dynamics. This aggregation bias is maximal close to the swamping limit of large-effect mutations and declines with increasing size of habitat 2, as predicted. Because of this aggregation bias, the contribution of large effect mutations to the phenotype *Q*_*A,Lap*_ given by equation (18) is largely overestimated.

**Figure 5.**
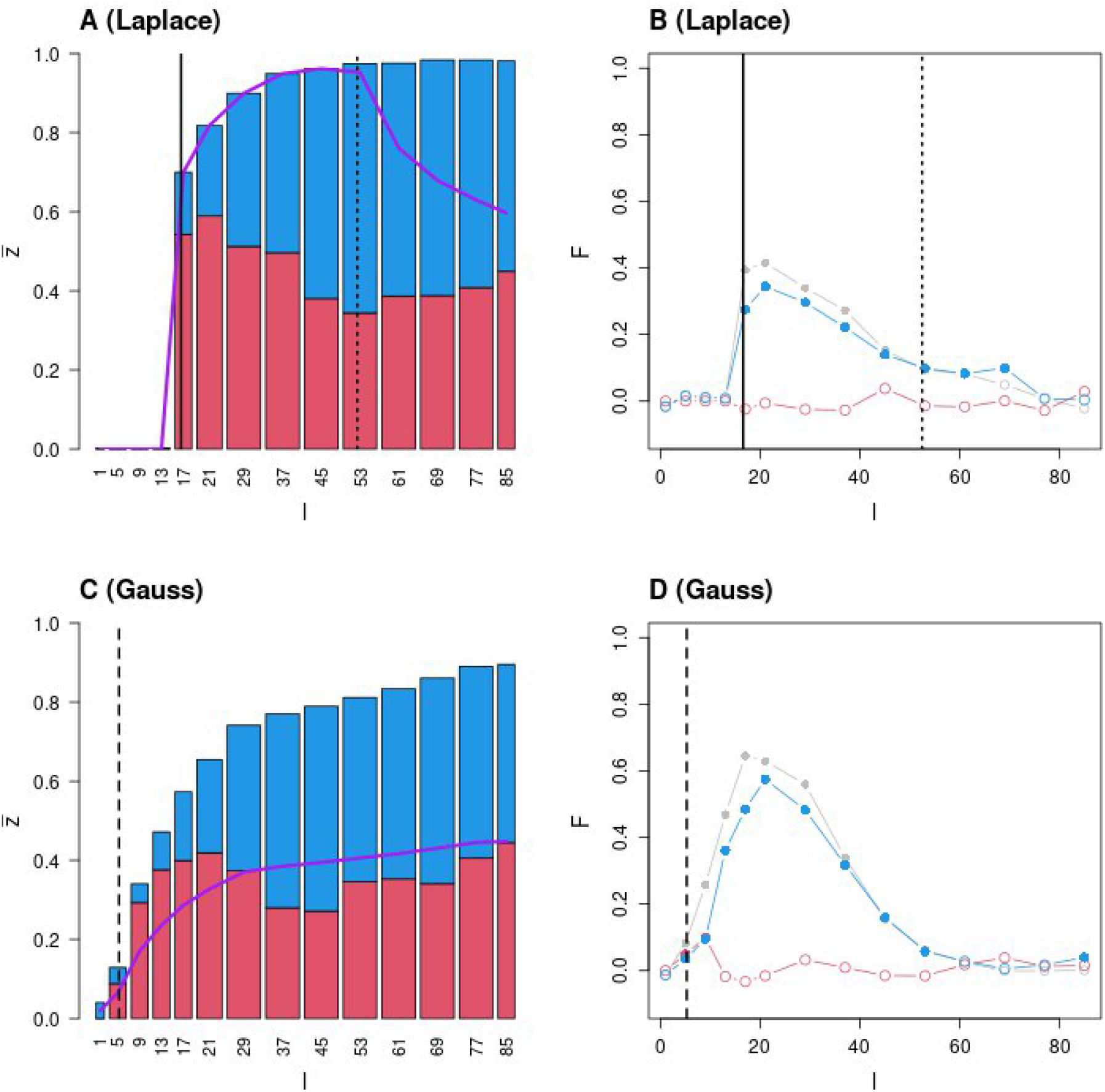
Short-term genetic architecture of local adaptation after the emergence phase. Composition and structure of adaptive genotypes at the center of habitat 2 after the initial phase of adaptation (see methods) for Laplace (panels A and B) and Gauss (panels C and D) fitness trade-off. Panels A and C present the average phenotype (bars) at the middle of habitat 2 for various sizes of habitat 2 (*l*), averaged over 20 replicates. The average relative contribution of alleles with large and small effects are presented in red and blue within each bar, respectively. The purple line shows the expected contribution of large versus small effects computed when rare at emergence (see eq. 17 and 18 of main text). In panel A and B (Laplace fitness function), vertical black lines show the analytical invasion thresholds above which a small (dotted line) or a large (solid line) effect mutation can invade alone the *z*\*=0 resident phenotype (*l*_*L*_(*a*) and *l*_*L*_(*A*) in main text, respectively). In panel C and D (Gauss fitness function), the vertical black dashed line show the analytical invasion threshold above which a small or large mutation can invade alone the *z*\*=0 resident phenotype (*l*_*G*_ in main text). Panels B and D present the average aggregation of adaptive alleles along the genetic map at the middle of habitat 2 for various sizes of habitat 2 (*l*), averaged over 20 replicates. Aggregation of all the adaptive alleles, small effects only and large effects only are presented in grey, blue and red, respectively. Filled dots correspond to significant F values, based on Bonferroni – corrected tests with 5% family-wise error rate.

#### Gauss trade-off

In the simulations shown on figure 5, the theoretical swamping limit of both small and large effect mutations (eq. 13) is *l*_*G*_ = 5.2. However, we do not observe a clear swamping effect in practice. Contrary to the Laplace case, the average trait value start to increase smoothly below this threshold (fig. 5C). With the Gauss fitness function, mutations initially enjoy an advantage in habitat 2, while suffering very limited fitness reduction in habitat 1. Hence, many alleles can persist long enough to generate a weak pattern of local adaptation. The contribution of large effect mutations predicted from equation (17) globally agrees with the simulation results, with moderate deviations. The “background trait value” bias (which favor large effect mutations) tend to cancel the “aggregation” bias (which favor small effect mutations). The aggregation bias remains noticeable for intermediate sizes of habitat 2 above the swamping limit (fig. 5D). Figure 7B shows a typical example of this dynamics.

### Simulation of the long-term reconfiguration phase

#### Laplace trade-off

Our prediction of swamping limit for large effect mutations *l*_*L*_*(A)* remains accurate in long-term simulations (fig. 6A). We do not observe a marked change in the contribution of small vs large effects to the adaptive phenotype compared to the initial phase. There is still an excess of large effect alleles compared to the control (where habitat 1 is absent), especially for sizes of habitat 2 close to the swamping limits. These large effect mutations do not show a significant pattern of aggregation at the end of this long-term phase (fig. 6B). In contrast, the aggregation of small effect mutations, which was already present in the initial phase, reinforces in the long term and reaches extreme values. This strong aggregation patterns occur for all sizes of habitat 2, even if it tends, like in the initial phase, to be stronger for sizes of habitat 2 close to the swamping limits.

**Figure 6.**
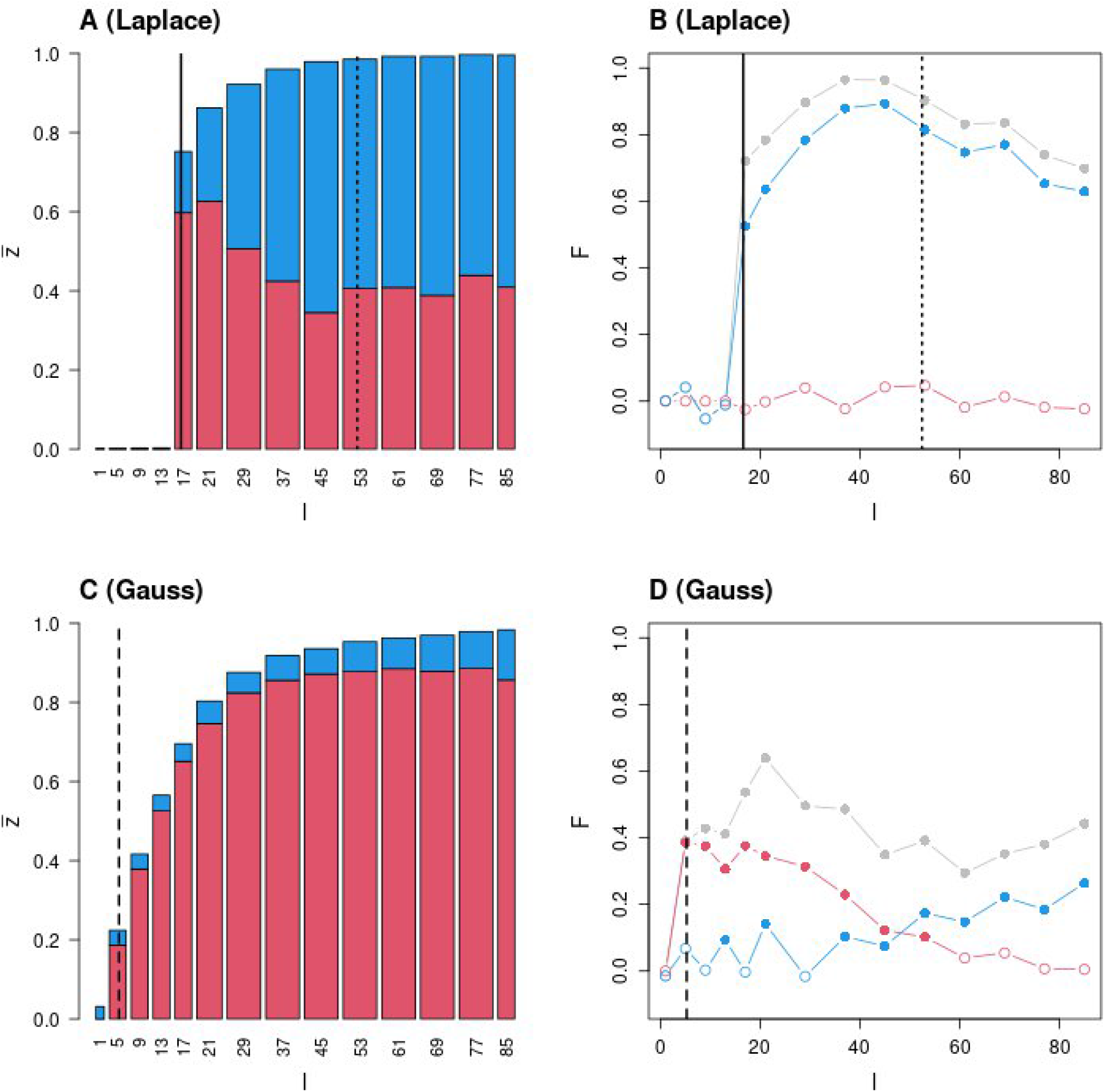
Long-term genetic architecture of local adaptation after the reconfiguration phase. The legend of the figure is identical than on fig. 5, except that measures are taken on the long term, after 400 000 generations.

**Figure 7.**
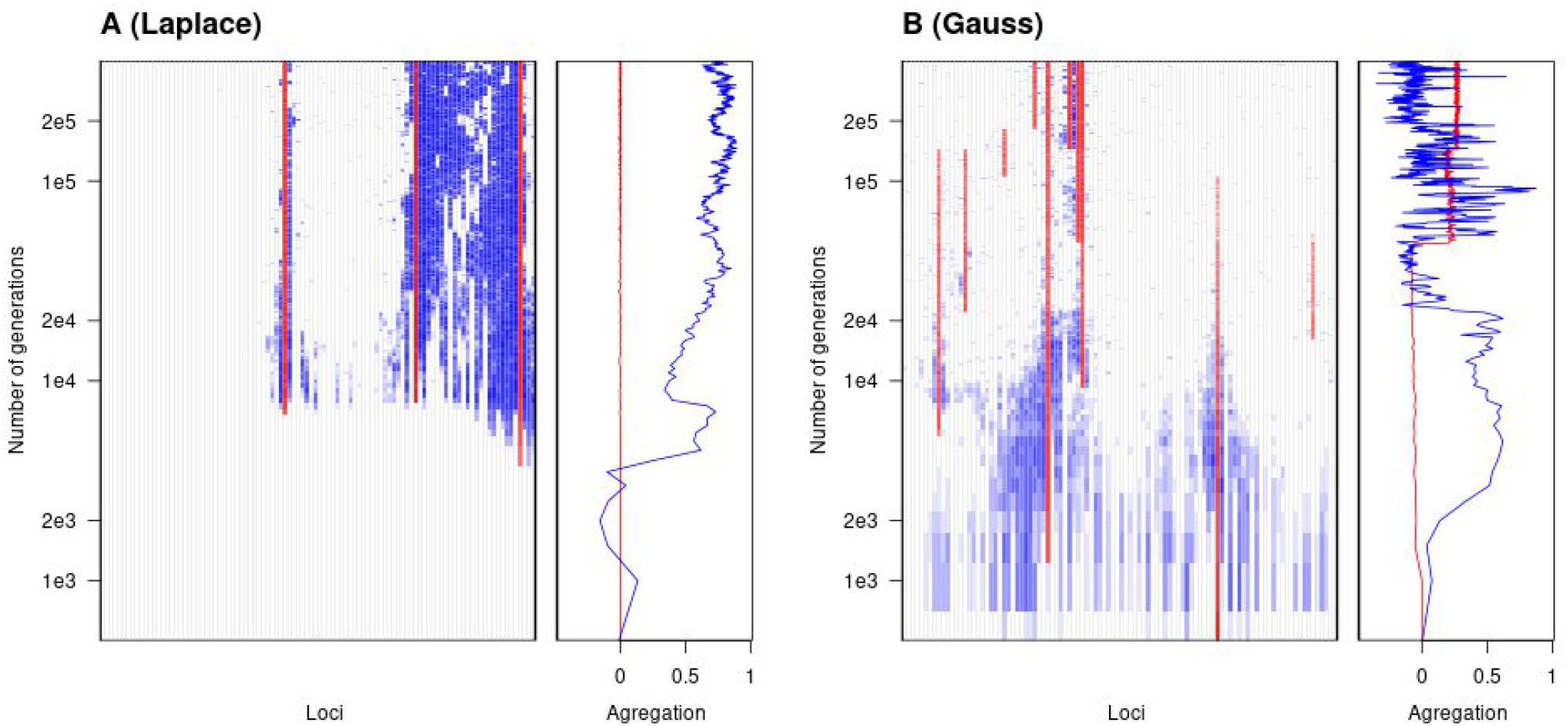
Typical examples of aggregation dynamics with Laplace (panel A) and Gauss (panel B) fitness functions. In each panel, the frequency of the dominant adaptive allele at each locus within the central deme of habitat 2 is presented, in blue if it is an allele with a small effect size, red if it is a large effect size, with darker color for larger frequency. The x-axis “loci” represent loci, as they are ordered on the chromosome The evolution of the genetic architecture through time can be observed along the y-axis (in log scale, which allows presenting simultaneously the emergence and long term reconfiguration phases). Right next to these maps, we present how the aggregation metrics of small (blue lines) and large effect size alleles (red shapes) change in time. In the two examples, the width of habitat 2 is set to *l*=19 demes (which is between the swamping limits for small and large effect mutations in the Laplace case, see fig. 5A).

#### Gauss trade-off

We observe weak local adaptation for sizes of habitat 2 below the swamping limit, with similar magnitude than during the initial phase of adaptation (fig. 6C). For all the habitat 2 sizes above the swamping limit, the contribution of large alleles to the phenotypic change markedly increases compared to the initial phase of adaptation. On the long run, small effects mutations are replaced by large effect mutations. For intermediate sizes of habitat above the swamping limit, these large effect mutations become aggregated (fig. 6D). For larger sizes of habitat 2, this aggregation of large effect mutations disappears, and is replaced by a pattern of aggregation of the residual small effect alleles. These residual small effect mutations strongly cluster around large effect mutations (the combined measure of aggregation on small and large effect mutations is much larger than each taken separately, fig. 6D).

## Discussion

### Local adaptation and aggregation in spatially explicit models

In spatially explicit models with limited dispersal, polymorphism can be maintained for a broad range of parameter values. Provided the environment is “coarse-grained” compared to the scale of gene flow (i.e. the sizes of the different habitats is sufficiently large), habitats’ cores are more preserved from gene flow compared to peripheral zones and clines form at the boundary between habitats. This phenomenon is not present in spatially implicit (two-patch or continent-island) models of adaptation. Here, for a given migration rate, local adaptation to a given habitat is always possible if the spatial extension of this habitat is large enough. Of course, a direct one-to-one comparison is not possible between spatially implicit and spatially explicit models, since the spatially explicit model introduces the effect of distance and extra-parameters. However, the spatially explicit model does not lead to the qualitative appreciation that polymorphism and aggregation are restricted to a small range of migration rates. As long as clines form at the boundary between habitats, selection tends to decrease recombination among alleles forming these clines in the patches close to the habitats boundary. Away from these clines, recombination is neutral since no polymorphism is present. Hence, these clines provide a stable context favoring recombination suppression (or equivalently aggregated architecture) in the long-term. Our work shows that an aggregated genetic architecture readily builds up and persist on the long term in these conditions, both through the initial recruitment of linked alleles (the “emergence” of local adaptation), and through the long-term competition between differentially aggregated genetic architectures (the “reconfiguration” phase).

### Condition for the existence of clines

The conditions of polymorphism in a spatially explicit model have been precisely worked out for a single locus using a diffusion approximation (Nagylaki 1975). In this population genetic model, the fitness effect of alleles in the different habitats are free parameters. The effect of the fitness trade-off (the ratio of selection coefficients outside versus inside the focal habitat) can be distinguished from the strength of selection (the selection coefficient within the focal habitat): the effect of each parameter can be kept constant while the other one is varying. In this context, and for a given fixed trade-off, the conditions for the existence of cline differ between weakly and strongly selected alleles. Alleles of large effect can more easily escape swamping and form clines.

Here, we adopt a quantitative genetic perspective and rather consider stabilizing selection on a trait with different optima in the different habitats. Selection is therefore determined by the difference in optimal phenotype across habitats, the intensity of stabilizing selection around each of these optima and the precise shape of the fitness function around them. The treatment of such models remains similar to population genetic models regarding the existence of a first cline at a single locus, without the complication of linkage and indirect selection (fig. 4). By using a fitness function relating the trait value to the fitness, it is possible to combine the effect of several loci in a parsimonious way. However, with stabilizing selection, the fitness effect of an allele with a given phenotypic effect depends on other alleles, contrary to simple population genetic models. For instance, if the population is at the optimum trait value, a mutation increasing the trait value is deleterious. It would be beneficial if the population had a trait value well below the optimal value. In addition to this background effect, the precise shape of the fitness function also matters. There are indeed many possible functions that can be used to describe stabilizing selection. It is necessary to scale them in some ways to compare results with different functions. A natural way to scale them is to assume that the overall fitness difference between habitats is kept constant (i.e., controlling the fitness drop of having, in one habitat, the phenotype that is optimal in the other). This scaling necessarily entails that alleles with a given phenotypic effect on the trait do not necessarily have the same fitness effect with the different functions, for a given average trait value in the population. The choice of the fitness function has a large qualitative impact on the condition of polymorphism, because it affects how the fitness trade-off of mutations (the ratio of selection coefficients across habitats) varies with the current phenotype and the size of the mutation phenotypic effect (fig. 2). In the Laplace fitness function, the trade-off is constant for alleles of small and large phenotypic effects, as long as the phenotypic effect does not overshoot the optimum, and the results are therefore very similar to the population genetic model where selection coefficients of alleles are constant parameters. There are conditions (of migration intensity and habitat size) where large effect alleles can form clines while small effect alleles cannot (fig. 4). In the Gauss fitness function, however, the fitness trade-off varies with the phenotypic effect of alleles and the intensity of selection. Intuitively, we see that when the population mean phenotype is close to the optimal value outside the pocket, an allele of small effect has an advantage in the pocket, but very little deleterious effect outside (since the fitness function is flat near the optimum outside the pocket). It can therefore invade very easily. For an allele with a larger effect, the trade-off is less favorable (it starts having a deleterious effect outside the pocket), but it enjoys a stronger intensity of selection. These two effects cancel each other, with the consequence that it is an all-or-nothing phenomenon: either all alleles can form clines or none. This Gauss case is equivalent to a population genetic model where the trade-off would vary with the intensity of selection (fig. 4).

Overall, the choice of the fitness function has a qualitative effect on the dynamic of local adaptation, especially regarding the relative contribution of alleles of small and large effects and their temporal dynamics. Previous models have considered various fitness function but have not explicitly shown that the choice of the fitness function had a qualitative effect on patterns of polygenic local adaptation when alleles of different phenotypic effects are considered (see section “Patterns of aggregation with different fitness functions” below for details and references).

### Condition to switch towards more aggregated architecture

Once local adaptation is established, aggregation can arise by the replacement of a given architecture by another (a change in the effect size distribution of alleles and/or in their positions along the chromosome). This progress towards aggregation takes time because the transition between architectures involves crossing a fitness valley. Indeed, when the phenotype is near its optimal value in the focal habitat, introducing a new local adaptation allele is likely to be deleterious, as it will cause an overshoot in the mean trait value. Hence, the first step for switching from one configuration to another involves recruiting an allele that has a deleterious effect in the context in which it arises. Of course, this deleterious effect can be immediately compensated by a frequency change at another locus (to keep the phenotype close to its optimum value), but this is necessarily a two-step phenomenon, where the first step is not favorable. Here too, the difference between the fitness functions affects the dynamics of the reconfiguration process. The Gauss fitness function is flat near the optimum, so that small deviations necessarily have small deleterious fitness effects in the focal habitat. Hence the fitness valley is shallower than in the Laplace case. As a consequence, transition between architecture is more fluid and occurs at a much faster rate in the Gauss compared to the Laplace case. In the latter, transitions involving large effect mutations are extremely difficult, and these alleles tend to be stable over very long period of time, even though small effect mutations can switch positions and end up forming large cluster around them. Again, this comparison is made scaling the overall fitness difference among habitats (as common in quantitative genetics models), and not the fitness effect of individual alleles (as common in population genetics models). When selection acts on a trait, the fitness effect of alleles depends on the context (the background trait value), and are therefore not directly scalable.

### Patterns of aggregation with different fitness functions

The vast majority of models of selection involving an environmental shift use Gauss fitness functions. This is the case for instance when studying a temporal environmental variation (e.g. Bürger and Lynch 1995; Gomulkiewicz and Houle 2009; Chevin et al. 2010; Matuszewski et al. 2015; Marshall et al. 2016; Anciaux et al. 2018). Spatial models of species range also most often use a Gauss fitness function (e.g. Kirkpatrick and Barton 1997; Alleaume-Benharira et al. 2006; Bridle et al. 2010; Polechová and Barton 2015; Fouqueau and Roze 2021). Such fitness function have received various justification and validation (Lande 1976; Martin and Lenormand 2006a; Martin et al. 2007). This fitness function is most often used for convenience for maintaining a normal distribution of phenotypes within a population, or to model smooth stabilizing selection around an optimum. As explained above, the choice of a fitness function also implicitly determines the fitness effects of mutations across environments and the shape of fitness trade-off across environments (Martin and Lenormand 2006b, 2015). While comparative analysis tend to globally support Gauss trade-off patterns around optima (Martin and Lenormand 2006b), it is difficult to empirically affirm that such trade-off are valid for all loci and alleles, especially for the small effect mutations that are difficult to study. The precise shape of fitness function has been shown to also be important in other circumstances. For instance, the shape of fitness functions away from optima is crucial for predicting demographic consequences of large departures from optimal phenotype in situation of rapid environmental change (Osmond and Klausmeier 2017). In any case, our results show that the dynamics and pattern of aggregation strongly depend on the fitness function used.

Although the two fitness functions considered here readily lead to aggregation patterns, the underlying configurations of alleles of small and large effects, their dynamics, and the resulting genetic map qualitatively differs. These qualitative differences between the Gauss and Laplace fitness function are caused by the conditions of polymorphism that differ for alleles of small and large effect, as explained above. This effect is particularly strong initially, when alleles are recruited to establish local adaptation. Here the crucial point is whether the fitness function is flat or not around the optimum outside the pocket, which determines whether small effect alleles have initially a more favorable trade-off than large effect alleles. The other important dynamical difference occurs in the longer term and depends on the ease to switch among architectures. Here, the crucial point is whether the fitness function is flat or not around the optimum inside the pocket, such that alternative architectures can evolve without having to cross a deep fitness valley.

Overall, with a Laplace trade-off, aggregation stems from the fact that small alleles tend to aggregate along few, non-aggregated large alleles, hence creating several small, but stable, chromosomal islands of adaptation around each of those large alleles. This effect is exacerbated when the pocket size is above the swamping limit of large effect alleles but below the swamping limit of small effect alleles. In this case, clines of small effects alleles emerge right from the beginning, as long as these alleles occur near large effect ones, and enjoy strong indirect selection. During this emergence phase, the interplay between the probability of invasion of small and large alleles and the hitch-hiking effects lead to a (relatively minor) non-linear change in the relative proportion of small vs. large effect alleles as the size of the habitat increases (fig. 5A). Because reconfiguration is slow with a Laplace fitness function, this effect is still present in the long term (fig. 6A). Overall, reconfiguration of the genetic architecture is markedly slower than in the Gauss case, and particularly rare regarding alleles with large effect sizes. As a result, the genetic architecture remains remarkably stable in time under this regime.

With a Gauss trade-off, all the alleles tend to aggregate together irrespectively of their effect size, while concentration (i.e. replacement of several small alleles by a large one) reinforces in time. This often leads to a single chromosomal island of adaptation in the long-term. Because of this concentration, effect sizes of adaptive alleles within the pocket are markedly higher with a Gauss than with a Laplace fitness trade-off in the long term (fig. 6A, C). Like for the Laplace case, there is initially a (relatively minor) non-linear change in the relative proportion of small vs. large effect alleles as the size of the habitat increases (fig. 5C, Appendix C). However, this effect disappears in the long term since reconfiguration occurs quite rapidly (fig. 6C).

Empirically, if the resolution of the genetic map is low, small chromosomic islands mentioned above may appear as single loci and aggregation may be overlooked. Therefore, chromosomal islands of adaptation may be more conspicuous under the Gauss trade-off, unless the concentration process allows for stacking all the necessary phenotypic effect on a single allele (Yeaman and Whitlock 2011). In contrast, under the Laplace trade-off, the local adaptation alleles may appear as scattered QTLs with large effect sizes. With a high resolution, however, alleles of small effects may be found surrounding these large effects loci. These expectations of course neglect that the architecture could also evolve because of chromosomal rearrangements (e.g. inversions) or local modifications of recombination rates (Lenormand and Otto 2000; Kirkpatrick and Barton 2006; Bürger and Akerman 2011; Lenormand 2012; Yeaman 2013; Charlesworth and Barton 2018). The importance of these secondary modifications of recombination ultimately depend on their relative rate of occurrence compared to the rate of architecture reconfiguration that we describe. In particular, they may play a stronger role in the Laplace case, where large effect alleles are stable on the long term and fail to consistently aggregate. However, these relative dynamics would require to be specifically investigated.

### Comparison to previous studies

In addition to the differences associated with considering a spatially explicit model with distance limited dispersal, there are several points to discuss in comparison to previous studies on local adaptation and genetic aggregation. First, all studies do not consider both the emergence and the reconfiguration phases in the process of aggregation, whereas the long term outcome depends on both phases. Second, all studies do not consider a fitness function relating trait to fitness, considering e.g. a set of loci with mutations with a constant effect. Stabilizing selection on a trait introduces the complication that the effect of alleles depends on the background and vary as local adaptation emerges. Third, models of local adaptation do not necessarily compare different shapes of fitness trade-off, or do so using cases that are apparently different, yet similar where it matters most. For instance Débarre and Gandon (2010) explored a wide range of trade-off functions in a two patch model. However, these functions are all similar on a critical aspect : their derivative of log-fitness close to optima is always 0. In that sense, all the trade-offs considered in that study are comparable to the Gauss trade-off here, which explains why they did not find a strong effect of the shape of the fitness trade-off on the pattern of local adaptation. Last, and this is a related point, the different studies have not systematically explored the effect of different fitness functions, even when they considered a flexible model allowing for such comparison. For instance, Yeaman and Whitlock (2011) considered a fitness function similar to ours in a two-patch model, with a flexible parameter allowing to explore Laplace and Gauss trade-offs considered in our study. However, the effect of this parameter is not clearly discussed in their study.

### Conclusion and perspective

We show that the process of genetic aggregation is a robust phenomenon when local adaptation evolves in a new environmental pocket. The detailed pattern of aggregation depends on both the emergence and the reconfiguration phase and vary depending on the precise shape of function relating trait values to fitness. The emergence of aggregation requires that several clines co-occur at the same spatial location. This situation may not systematically characterize ecological variation. For instance, it is possible that local adaptation occurs on ecological gradients (notably abiotic such as temperature, photoperiod etc.), rather than on “pockets” of habitats. In this case, staggered clines may evolve with little scope for genetic aggregation. However, the presence of an abiotic gradient does not necessarily translate into a gradient in optimal trait values in a given species. We have a limited knowledge of this variation. Moreover, many natural habitats may be better described as patches of different sizes rather than as gradients. In particular, anthropized landscapes may be characterized by the creation of marked segmentation and abrupt transitions in habitat quality at moderate-to-coarse spatial grains (typically 0.1-1000 ha, Strayer 2005) due to infrastructures (e.g. roads), spatial heterogeneity in land use (e.g. crops in agricultural landscapes or stands within managed forests) and management practices (e.g. pesticide application). This first trend may combine with a homogenization process at fine grain (typically <0.1 ha; Strayer 2005) in many contexts like urban areas (e.g. soil artificialization) or managed forest or agricultural ecosystems (e.g. Fraterrigo et al. 2005 and references therein). Hence, the combination of both processes suggests that the rapidly growing human imprint on landscape structure should often result in a marked patchiness, making spatially explicit models essential to investigate patterns and dynamics of local adaptation in the field, and more generally adaptation and resilience of organisms to global changes.

## Supporting information

AppendicesABC

## Acknowledgements

We thank Noémie Harmand and Florence Débarre for discussions and comments. We also thank PCI reviewers, recommender (C. Mullon) and managers.

## References

Alleaume-Benharira, M., I. R. Pen, and O. Ronce. 2006. Geographical patterns of adaptation within a species’ range: interactions between drift and gene flow. J. Evol. Biol. 19:203–215.

Anciaux, Y., L. M. Chevin, O. Ronce, and G. Martin. 2018. Evolutionary rescue over a fitness landscape. Genetics 209:265–279.

Barton, N. H., and G. M. Hewitt. 1989. Adaptation, speciation and hybrid zones. Nature 341:497–503.

Bridle, J. R., J. Polechova, M. Kawata, and R. K. Butlin. 2010. Why is adaptation prevented at ecological margins? New insights from individual-based simulations. Ecol. Lett. 13:485–494.

Bürger, R., and A. Akerman. 2011. The effects of linkage and gene flow on local adaptation: A two-locus continent-island model. Theor. Popul. Biol. 80:272–288. Elsevier Inc.

Bürger, R., and M. Lynch. 1995. Evolution and Extinction in a Changing Environment: A Quantitative-Genetic Analysis. Evolution 49:151–163.

Charlesworth, B., and N. H. Barton. 2018. The spread of an inversion with migration and selection. Genetics 208:377–382.

Charlesworth, D. 2016. The status of supergenes in the 21st century: Recombination suppression in Batesian mimicry and sex chromosomes and other complex adaptations. Evol. Appl. 9:74–90.

Chevin, L. M., R. Lande, and G. M. Mace. 2010. Adaptation, Plasticity, and Extinction in a Changing Environment: Towards a Predictive Theory. PLoS Biol. 8:e1000357.

Débarre, F., and S. Gandon. 2010. Evolution of specialization in a spatially continuous environment. J. Evol. Biol. 23:1090–1099.

Flaxman, S. M., J. L. Feder, and P. Nosil. 2013. Genetic hitchhiking and the dynamic buildup of genomic divergence during speciation with gene flow. Evolution 67:2577–2591.

Fouqueau, L., and D. Roze. 2021. The evolution of sex along an environmental gradient. Evolution 75:1334–1347.

Fraterrigo, J. M., M. G. Turner, S. M. Pearson, and P. Dixon. 2005. Effects of past land use on spatial heterogeneity of soil nutrients in southern appalachian forests. Ecol. Monogr. 75:215–230.

Gomulkiewicz, R., and D. Houle. 2009. Demographic and genetic constraints on evolution. Am. Nat. 174.

Haldane, J. (1927). A mathematical theory of natural and artificial selection, part V: selection and mutation. Mathematical Proceedings of the Cambridge Philosophical Society 23(7):838–844.

Jonsson, M. 2003. Colonisation ability of the threatened tenebrionid beetle Oplocephala haemorrhoidalis and its common relative Bolitophagus reticulatus. Ecological Entomology 28: 159–167.

Karlin, S. 1976. Population subdivision and selection migration interaction. BT - Population genetics and ecology Pp. 616–657 in S. Karlin and E. Nevo, eds. Academic Press Inc., New York, NY.

Kirkpatrick, M., and N. Barton. 2006. Chromosome inversions, local adaptation and speciation. Genetics 173:419–34.

Kirkpatrick, M., and N. H. Barton. 1997. Evolution of a species’ range. Am. Nat. 150:1–23.

Lande, R. 1976. Natural-Selection and Random Genetic Drift in Phenotypic Evolution. Evolution 30:314–334.

Lenormand, T. 2012. From local adaptation to speciation: specialization and reinforcement. Int. J. Ecol. 2012:e508458.

Lenormand, T. 2002. Gene flow and the limits to natural selection. Trends Ecol. Evol. 17:183–189.

Lenormand, T. 2003. The evolution of sex dimorphism in recombination. Genetics 163:811–822.

Lenormand, T., and S. P. P. Otto. 2000. The evolution of recombination in a heterogeneous environment. Genetics 156:423–438.

Marshall, D. J., S. C. Burgess, and T. Connallon. 2016. Global change, life-history complexity and the potential for evolutionary rescue. Evol. Appl. 9:1189–1201.

Martin, G., S. F. S. F. Elena, and T. Lenormand. 2007. Distributions of epistasis in microbes fit predictions from a fitness landscape model. Nat. Genet. 39:555–560.

Martin, G., and T. Lenormand. 2006a. A general multivariate extension of Fisher’s geometrical model and the distribution of mutation fitness effects across species. Evolution 60:893–907.

Martin, G., and T. Lenormand. 2006b. The fitness effect of mutations across environments: a survey in the light of fitness landscape models. Evolution 60:2413–2427.

Martin, G., and T. Lenormand. 2015. The fitness effect of mutations across environments: Fisher’s geometrical model with multiple optima. Evolution 69:1433–1447.

Matuszewski, S., J. Hermisson, and M. Kopp. 2015. Catch me if you can: Adaptation from standing genetic variation to a moving phenotypic optimum. Genetics 200:1255–1274.

Nagylaki, T. 1975. Conditions for the existence of clines. Genetics 80:595–615.

Nosil, P., D. J. Funk, and D. Ortiz-Barrientos. 2009. Divergent selection and heterogeneous genomic divergence. Mol. Ecol. 18:375–402.

Osmond, M. M., and C. A. Klausmeier. 2017. An evolutionary tipping point in a changing environment. Evolution 71:2930–2941.

Otto, S. P. 2009. The Evolutionary Enigma of Sex. Am. Nat. 174:S1–S14.

Otto, S. P. S. P., and T. Lenormand. 2002. Resolving the paradox of sex and recombination. Nat. Rev. Genet. 3:252–261.

Polechová, J., and N. H. Barton. 2015. Limits to adaptation along environmental gradients. Proc. Natl. Acad. Sci. U. S. A. 112:6401–6406.

Pylkov, K. V, L. A. Zhivotovsky, and M. W. Feldman. 1998. Migration versus mutation in the evolution of recombination under multilocus selection. Genet. Res. 71:247–256.

Ravigne, V., U. Dieckmann, and I. Olivieri. 2009. Live where you thrive: joint evolution of habitat choice and local adaptation facilitates specialization and promotes diversity. Am. Nat. 174:E141–E169.

Sardell, J. M., and M. Kirkpatrick. 2020. Sex differences in the recombination landscape. Am. Nat. 195:361–379.

Schwander, T., R. Libbrecht, and L. Keller. 2014. Supergenes and Complex Phenotypes. Curr. Biol. 24:R288–R294.

Slatkin, M. 1973. Gene flow and selection in a cline. Genetics 75:733–756.

Slatkin, M. 1978. Spatial patterns in the distributions of polygenic characters. J. Theor. Biol. 70:213–228.

Strasburg, J. L., N. A. Sherman, K. M. Wright, L. C. Moyle, J. H. Willis, and L. H. Rieseberg. 2012. What can patterns of differentiation across plant genomes tell us about adaptation and speciation? Philos. Trans. R. Soc. B Biol. Sci. 367:364–373.

Strayer, D. L. 2005. Challenges in Understanding the Functions of Ecological Heterogeneity BT - Ecosystem Function in Heterogeneous Landscapes. Pp. 411–425 in G. M. Lovett, M. G. Turner, C. G. Jones, and K. C. Weathers, eds. Springer New York, New York, NY.

Yeaman, S. 2013. Genomic rearrangements and the evolution of clusters of locally adaptive loci. Proc. Natl. Acad. Sci. U. S. A. 110:E1743–E1751.

Yeaman, S., S. Aeschbacher, and R. Bürger. 2016. The evolution of genomic islands by increased establishment probability of linked alleles. Mol. Ecol. 25:2542–2558.

Yeaman, S., and M. C. Whitlock. 2011. The genetic architecture of adaptation under migration-selection balance. Evolution 65:1897–1911.

