## AppendicesABC for "The genetic architecture of local adaptation in a cline"

### Appendix A : Invasion fitness of a small mutant

We considered one adaptive locus with two alleles 1 and 2. We note $w_{iab}$ the fitness of an individual with one copy of alleles $a$ and one copy of allele $b$ within site $i$. The frequency of allele 2 in site $i$, denoted $x_i$ follow the following dynamics:

$$x_{i2}(t+1) = \frac{\sum_{j=1}^{n} \pi_{ij}\left[w_{i12}x_{j2}(t)(1-x_{j2}(t))+w_{i22}x_{j2}(t)^2\right]}{\sum_{j=1}^{n} \pi_{ij}\left[w_{i11}(1-x_{j2}(t))^2+2w_{i12}x_{j2}(t)(1-x_{j2}(t))+w_{i22}x_{j2}(t)^2\right]}$$

When allele 2 is rare, the above dynamics becomes:

$$x_{i2}(t+1) = \frac{w_{i12}}{w_{i11}} \sum_{j=1}^{n} \pi_{ij} x_{j2}(t)$$

which can be written in a classic matricial form for rare mutant invasion in a spatialized system under soft-selection regime (Karlin, 1976):

$$\boldsymbol{x_2}(t+1) = \boldsymbol{D\Pi x_2}(t) \tag{A1}$$

where $\boldsymbol{D}$ is a $n \times n$ diagonal matrix such that $D_{ii} = \frac{w_{i12}}{w_{i11}}$. Note that equation (A1) corresponds to equation (4) in the main text. Invasion is possible if and only if the dominant eigenvalue of $\boldsymbol{D\Pi}$ has a module strictly above 1. When the mutant has exactly the same fitness than the resident allele (i.e. $\forall i \in \{1, 2, ..., n\}, w_{i12} = w_{i11}$), the leading eigenvalue of $\boldsymbol{D\Pi} = \boldsymbol{\Pi}$ is 1, and $(1, 1, ..., 1)$ is an associated eigenvector. We now assume that the mutant has a slightly different fitness in sites such that $\frac{w_{i12}}{w_{i11}} = 1 + s_i\epsilon$.

Biogically speaking, $s_i\epsilon$ is the selection coefficient of the mutant allele in site $i$, and $s_i$ is the corresponding selection gradient.

We look for the leading eigenvalue $\lambda$ of $\boldsymbol{D\Pi}$ under the form:

$$\lambda = 1 + \lambda_1\epsilon + \lambda_2\epsilon^2 + o(\epsilon^2)$$

and an associated eigenvector $\boldsymbol{v}$ of the form:

$$\boldsymbol{v} = \left(1 + v_{11}\epsilon + v_{12}\epsilon^2 + o(\epsilon^2), ..., 1 + v_{n1}\epsilon + v_{n2}\epsilon^2 + o(\epsilon^2)\right)$$

Equation (A1) then implies that for any site $i$:

$$\begin{pmatrix} 1 \\ +(\lambda_1 + v_{i1})\,\epsilon \\ +(v_{i2} + \lambda_1 v_{i1} + \lambda_2)\,\epsilon^2 \\ +o\left(\epsilon^2\right) \end{pmatrix} = \begin{pmatrix} 1 \\ +\left(s_i + \sum_{j=1}^{n} \Pi_{ij} v_{j1}\right)\epsilon \\ +\left(s_i\left(\sum_{j=1}^{n} \Pi_{ij} v_{j1}\right) + \sum_{j=1}^{n} \Pi_{ij} v_{j2}\right)\epsilon^2 \\ +o\left(\epsilon^2\right) \end{pmatrix}$$

Identification of expansion terms yields that for all $i \in \{1, 2, ..., n\}$:

$$\lambda_1 + v_{i1} = s_i + \sum_{j=1}^{n} \Pi_{ij} v_{j1} \tag{A2a}$$

$$v_{i2} + \lambda_1 v_{i1} + \lambda_2 = s_i \left( \sum_{j=1}^{n} \Pi_{ij} v_{j1} \right) + \sum_{j=1}^{n} \Pi_{ij} v_{j2} \tag{A2b}$$

Summing the equation (A2a) over $i$:

$$\begin{aligned} & n\lambda_1 + \textstyle\sum_{i=1}^{n} v_{i1} = \sum_{i=1}^{n} s_i + \sum_{i=1}^{n} \sum_{j=1}^{n} \Pi_{ij} v_{j1} \\ \iff & n\lambda_1 + \textstyle\sum_{i=1}^{n} v_{i1} = \sum_{i=1}^{n} s_i + \sum_{j=1}^{n} (\sum_{i=1}^{n} \Pi_{ij}) v_{j1} \\ \iff & n\lambda_1 + \textstyle\sum_{i=1}^{n} v_{i1} = \sum_{i=1}^{n} s_i + \sum_{j=1}^{n} v_{j1} \\ \iff & n\lambda_1 = \textstyle\sum_{i=1}^{n} s_i \end{aligned}$$

and we obtain:

$$\lambda_1 = \frac{1}{n} \sum_{i=1}^{n} s_i = \bar{s} \tag{A3}$$

which entails that for all $i \in \{1, 2, ..., n\}$:

$$v_{i1} - \sum_{j=1}^{n} \Pi_{ij} v_{j1} = s_i - \bar{s}$$

This can be summarized in the matrix equality:

$$(\boldsymbol{I} - \boldsymbol{\Pi})\, \boldsymbol{v_1} = \boldsymbol{s_0}$$

where $\boldsymbol{I}$ is the identity matrix of dimension $n$, $\boldsymbol{v_1} = (v_{11}, v_{21}, ..., v_{n1})$ and $\boldsymbol{s_0}$ is the vector of centered relative fitnesses of the mutant across the landscape: $s_{0i} = s_i - \bar{s}$. Previous equality implies that, formally :

$$\boldsymbol{v_1} = \sum_{k=0}^{+\infty} \boldsymbol{\Pi}^k \boldsymbol{s_0} \tag{A4}$$

One needs to check that the formal expansion (A4) is well defined. To do so, we use Fourier analysis as detailed e.g. in Rousset (2004). We assume that the number of sites $n$ is uneven and labelled in $\{-\frac{n-1}{2}, ..., \frac{n-1}{2}\}$. Define the family $(\boldsymbol{\omega_k})_{k \in \{-\frac{n-1}{2}, ..., \frac{n-1}{2}\}} \in \mathbb{C}^n$ such that the $l$ coordinate of $\boldsymbol{\omega_k}$ is

$\omega_{kl} = e^{\frac{i2\pi k}{n}(l-1)}$. ($\boldsymbol{\omega_k}$) is a basis of $\mathbb{C}^n$ viewed as a $\mathbb{C}$-vectorial space. Define $\sigma_k$, the coordinate of $\boldsymbol{s_0}$ along $\boldsymbol{\omega_k}$:

$$\boldsymbol{s_0} = \sum_{k=-\frac{n-1}{2}}^{\frac{n-1}{2}} \sigma_k \boldsymbol{\omega_k}$$

Assuming that $\boldsymbol{\Pi}$ is homogeneous and isotropic, it can be shown that $\boldsymbol{\Pi}\boldsymbol{\omega_k} = nc_{-k}\boldsymbol{\omega_k}$, where $c_k = c_{-k} \in \mathbb{R}$ is the coordinate of the first row of $\boldsymbol{\Pi}$ along $\boldsymbol{\omega_k}$. Therefore :

$$\begin{aligned} \boldsymbol{\Pi}^k \boldsymbol{s_0} &= \sum_{l=-\frac{n-1}{2}}^{\frac{n-1}{2}} \sigma_l \boldsymbol{\Pi}^k \boldsymbol{\omega_l} \\ &= \sum_{l=-\frac{n-1}{2}}^{\frac{n-1}{2}} \sigma_l \left(nc_{-l}\right)^k \boldsymbol{\omega_l} \\ &= 2\sum_{l=1}^{\frac{n-1}{2}} \sigma_l \left(nc_l\right)^k Re\left(\boldsymbol{\omega_l}\right) \end{aligned} \tag{A5}$$

where we used $\sigma_0 = 0$ because $\boldsymbol{s_0}$ is centered. In main text, we considered an isotropic and homogeneous kernel which takes the form : $\boldsymbol{\Pi} = (1-m)\boldsymbol{I} + m\boldsymbol{\Delta}$, where $\Delta_{ij} = 1_{d_{ij} \leq d}$. Then the $c_k$ can be computed for $k \in \{1, ..., \frac{n-1}{2}\}$ as:

$$nc_k = 1 - m + m\left(\frac{\frac{1}{2d+1}\sin\left(\frac{\pi k(2d+1)}{n}\right)}{\sin\left(\frac{\pi k}{n}\right)}\right) \tag{A6}$$

Using the concavity of $x \to \sin\left(\frac{\pi x}{n}\right)$ on $\left[0, \frac{n}{2}\right]$, one can show that $|nc_k| < 1$ for $k \in \{1, ..., \frac{n-1}{2}\}$, ensuring that the formal expansion (A4) of $\boldsymbol{v_1}$ is well defined.

Combining (A4) and (A5) yields:

$$\boldsymbol{v_1} = 2\sum_{l=1}^{\frac{n-1}{2}} \frac{\sigma_l}{1 - nc_l} Re\left(\boldsymbol{\omega_l}\right) \tag{A7}$$

Using $\lambda_1 = \bar{s}$, (A2b) yields for all $i \in \{1, 2, ..., n\}$:

$$v_{i2} + \bar{s}v_{i1} + \lambda_2 = s_i \sum_{j=1}^{n} \Pi_{ij} v_{j1} + \sum_{j=1}^{n} \Pi_{ij} v_{j2}$$

Summing over $i$:

$$\begin{aligned}
&\sum_{i=1}^{n} v_{i2} + \bar{s}\sum_{i=1}^{n} v_{i1} + n\lambda_2 = \sum_{i=1}^{n} s_i \sum_{j=1}^{n} \Pi_{ij} v_{j1} + \sum_{i=1}^{n}\sum_{j=1}^{n} \Pi_{ij} v_{j2} \\
&\iff \sum_{i=1}^{n} v_{i2} + \bar{s}\sum_{i=1}^{n} v_{i1} + n\lambda_2 = \sum_{i=1}^{n}\sum_{j=1}^{n} s_i \Pi_{ij} v_{j1} + \sum_{j=1}^{n} (\sum_{i=1}^{n} \Pi_{ij}) v_{j2} \\
&\iff \sum_{i=1}^{n} v_{i2} + \bar{s}\sum_{i=1}^{n} v_{i1} + n\lambda_2 = \sum_{i=1}^{n}\sum_{j=1}^{n} s_i \Pi_{ij} v_{j1} + \sum_{j=1}^{n} v_{j2} \\
&\iff \bar{s}\sum_{i=1}^{n} v_{i1} + n\lambda_2 = \sum_{i=1}^{n}\sum_{j=1}^{n} s_i \Pi_{ij} v_{j1} \\
&\iff n\lambda_2 = \sum_{i=1}^{n}\sum_{j=1}^{n} s_i \Pi_{ij} v_{j1} - \bar{s}\sum_{i=1}^{n} v_{i1}
\end{aligned}$$

and we obtain :

$$\lambda_2 = \frac{1}{n}\,{}^t\boldsymbol{s}\boldsymbol{\Pi}\boldsymbol{v_1} - \bar{s}\bar{v}_1$$

where $\bar{v}_1 = \frac{1}{n}\sum_{i=1}^{n} v_{i1}$. Using (A7) we obtain:

$$\begin{aligned}
\lambda_2 &= \tfrac{2}{n}\sum_{l=1}^{\frac{n-1}{2}} \tfrac{\sigma_l}{1-nc_l}\,{}^t\boldsymbol{s}\boldsymbol{\Pi} Re\left(\boldsymbol{\omega_l}\right) \\
&= \tfrac{1}{n}\sum_{l=1}^{\frac{n-1}{2}} \tfrac{\sigma_l}{1-nc_l}\,{}^t\boldsymbol{s}\left(\boldsymbol{\Pi}\boldsymbol{\omega_l} + \boldsymbol{\Pi}\boldsymbol{\omega_{-l}}\right) \\
&= \tfrac{1}{n}\sum_{l=1}^{\frac{n-1}{2}} \tfrac{\sigma_l}{1-nc_l}\,{}^t\boldsymbol{s}\left(nc_{-l}\boldsymbol{\omega_l} + nc_l\boldsymbol{\omega_{-l}}\right) \\
&= \sum_{l=1}^{\frac{n-1}{2}} \tfrac{\sigma_l c_l}{1-nc_l}\,{}^t\boldsymbol{s}\left(\boldsymbol{\omega_l} + \boldsymbol{\omega_{-l}}\right) \\
&= \sum_{l=1}^{\frac{n-1}{2}} \tfrac{\sigma_l c_l}{1-nc_l}\,{}^t\boldsymbol{s}\left(\boldsymbol{\omega_l} + \boldsymbol{\omega_{-l}}\right) \\
&= n\sum_{l=1}^{\frac{n-1}{2}} \tfrac{\sigma_l c_l}{1-nc_l}\left(\sigma_l + \sigma_{-l}\right) \\
&= 2n\sum_{l=1}^{\frac{n-1}{2}} \tfrac{\sigma_l^2 c_l}{1-nc_l}
\end{aligned} \tag{A8}$$

The environment is made of $l$ patches of type 2 and $n-l$ patches of type 1. $l$ is assumed uneven below. The $l$ contiguous sites of environment 2 can be positionned on the circle of sites from label $-\frac{l-1}{2}$ to label $\frac{l-1}{2}$ without loss of generality. We define the selection gradients $s_1, s_2 = \left(\frac{w_{i12}}{w_{i11}} - 1\right)/\epsilon$ for sites $i$ belonging to environment 1 and 2, respectively. Coordinates of the vector of centered relative fitnesses of the mutant across the landscape $\boldsymbol{s_0}$ are:

$$\begin{cases} s_{0i} = s_2 - \bar{s} \text{ if } -\frac{l-1}{2} \le i \le \frac{l-1}{2} \\ s_{0i} = s_1 - \bar{s} \text{ if } i \le -\frac{l-1}{2} - 1 \text{ or } i \ge \frac{l-1}{2} + 1 \end{cases}$$

Recalling that for $k \in \{1, ..., \frac{n-1}{2}\}$:

$$\sum_{l=-\frac{n-1}{2}}^{\frac{n-1}{2}} s_{0l}\bar{\omega}_{kl} = \sum_{l=-\frac{n-1}{2}}^{\frac{n-1}{2}} \sum_{m=-\frac{n-1}{2}}^{\frac{n-1}{2}} \sigma_m \omega_{ml}\bar{\omega}_{kl} = \sum_{m=-\frac{n-1}{2}}^{\frac{n-1}{2}} \sigma_m \sum_{l=-\frac{n-1}{2}}^{\frac{n-1}{2}} \omega_{ml}\bar{\omega}_{kl} = n\sigma_k$$

it holds that:

$$\begin{aligned}
&n\sigma_k = \sum_{l=-\frac{n-1}{2}}^{\frac{n-1}{2}} s_{0l}\bar{\omega}_{kl} \\
&= \sum_{l=-\frac{l-1}{2}}^{\frac{l-1}{2}} (s_2 - \bar{s})\bar{\omega}_{kl} + \sum_{l=-\frac{n-1}{2}}^{-\frac{l-1}{2}-1} (s_1 - \bar{s})\bar{\omega}_{kl} + \sum_{l=\frac{l-1}{2}+1}^{\frac{n-1}{2}} (s_1 - \bar{s})\bar{\omega}_{kl} \\
&= (s_2 - s_1) \sum_{l=-\frac{l-1}{2}}^{\frac{l-1}{2}} \bar{\omega}_{kl} + (s_1 - \bar{s}) \sum_{l=-\frac{n-1}{2}}^{\frac{n-1}{2}} \bar{\omega}_{kl} \\
&= (s_2 - s_1) \frac{1-e^{-\frac{i2\pi k}{n}l}}{1-e^{-\frac{i2\pi k}{n}}} e^{\frac{i2\pi k}{n}\frac{l-1}{2}} \\
&= (s_2 - s_1) \frac{\sin\left(\frac{\pi k l}{n}\right)}{\sin\left(\frac{\pi k}{n}\right)}
\end{aligned} \tag{A9}$$

Using (A9) in (A8) yields:

$$\lambda_2 = \frac{2}{n} (s_2 - s_1)^2 \sum_{q=1}^{\frac{n-1}{2}} \left( \frac{\sin\left(\frac{\pi q l}{n}\right)}{\sin\left(\frac{\pi q}{n}\right)} \right)^2 \frac{c_q}{1 - n c_q}$$

and using also (A6) yields:

$$\lambda_2 = \frac{2 (s_2 - s_1)^2}{m n^2} \sum_{q=1}^{\frac{n-1}{2}} \left( \frac{\sin\left(\frac{\pi q l}{n}\right)}{\sin\left(\frac{\pi q}{n}\right)} \right)^2 \frac{1 - m + m \left( \frac{\frac{1}{(2d+1)} \sin\left(\frac{\pi q (2d+1)}{n}\right)}{\sin\left(\frac{\pi q}{n}\right)} \right)}{1 - \frac{\frac{1}{(2d+1)} \sin\left(\frac{\pi q (2d+1)}{n}\right)}{\sin\left(\frac{\pi q}{n}\right)}} \tag{A10}$$

Taking the limit $n \to +\infty$ in (A3) and (A10), one obtains:

$$\begin{aligned}
\lambda_1 &\approx s_1 \\
\lambda_2 &\approx \frac{2(s_2-s_1)^2 l^2}{m} \sum_{q=1}^{+\infty} \frac{1}{\frac{1}{6}((2d+1)^2-1)(\pi q)^2} = \frac{2(s_2-s_1)^2 l^2}{m((2d+1)^2-1)}
\end{aligned}$$

Consequently, the expansion to order 2 of the dominant eigenvalue of $\boldsymbol{D\Pi}$ for a mutant with fitness variation $\epsilon \boldsymbol{s}$ in space is:

$$\lambda = 1 + s_1\epsilon + \frac{2\left(s_2\epsilon - s_1\epsilon\right)^2 l^2}{m\left((2d+1)^2 - 1\right)} + o(\epsilon^2) \tag{A11}$$

Developing fitness to order 2 with respect to the mutant allele phenotypic effect $a$ yields :

$$s_i\epsilon = \frac{w_i(z+a)}{w_i(z)} - 1 = \frac{w_i'(z)}{w_i(z)} a + \frac{w_i''(z)}{2w_i(z)} a^2 + o(a^2)$$

which we can use in (A11) to obtain an expansion of order 2 of mutant invasion fitness as a function of its phenotypic effect:

$$\lambda = 1 + \frac{w_1'(z)}{w_1(z)} a + \left[ \frac{1}{2} \frac{w_1''(z)}{w_1(z)} + \frac{2 \left( \frac{w_2'(z)}{w_2(z)} - \frac{w_1'(z)}{w_1(z)} \right)^2 l^2}{m \left( (2d+1)^2 - 1 \right)} \right] a^2 + o(a^2)$$

Using the variance of the dispersal kernel $\sigma^2 = d(d+1)/3$, we finally obtain:

$$\lambda = 1 + \frac{w_1'(z)}{w_1(z)} a + \left[ \frac{w_1''(z)}{w_1(z)} + \frac{\left( \frac{w_2'(z)}{w_2(z)} - \frac{w_1'(z)}{w_1(z)} \right)^2 l^2}{3m\sigma^2} \right] \frac{a^2}{2} + o(a^2)$$

which is the expression used in equations (6a) and (6b) of main text.

#### References

Karlin, S. (1976). Population subdivision and selection migration interaction. In *Population genetics and ecology*, pages 616–657. Elsevier Science, Saint Louis.

Rousset, F. (2004). Spatially homogeneous dispersal: the island model and isolation by distance. In *Genetic structure and selection in subdivided populations*, volume 40, pages 23–52. Princeton University Press.

### Appendix B : Computing fitness invasion criterion and associated quantities for each fitness function

#### General definitions and results in the main text

$$s_1(z) = w_1'(z)/w_1(z)$$
$$t_1(z) = w_1''(z)/w_1(z)$$
$$S_1(\epsilon) = w_1(z+\epsilon)/w_1(z) - 1$$
$$s_2(z) = w_2'(z)/w_2(z)$$
$$t_2(z) = w_2''(z)/w_2(z)$$
$$S_2(\epsilon) = w_2(z+\epsilon)/w_2(z) - 1$$

##### Nagylaki parameters

$$k = (l/\sqrt{2})\sqrt{S_2(\epsilon)}/(\sigma\sqrt{m}) \qquad \text{(eq. 7a of maintext)}$$
$$\alpha = -S_1(\epsilon)/S_2(\epsilon) \qquad \text{(eq. 7b of maintext)}$$

Expanding selection coefficients $S_1(\epsilon), S_2(\epsilon)$ to order 2 with respect to $\epsilon$ yields:

$$S_1(\epsilon) = s_1\epsilon + t_1\frac{\epsilon^2}{2} + o(\epsilon^2)$$
$$S_2(\epsilon) = s_2\epsilon + t_2\frac{\epsilon^2}{2} + o(\epsilon^2)$$

Therefore:

$$k = \frac{(l/\sqrt{2})\sqrt{s_2}}{\sigma\sqrt{m}}\sqrt{\epsilon} + o(\sqrt{\epsilon})$$

$$\begin{aligned}
\alpha &= -\frac{s_1+t_1\frac{\epsilon}{2}+o(\epsilon)}{s_2+t_2\frac{\epsilon}{2}+o(\epsilon)} \\
&= -\frac{s_1+t_1\frac{\epsilon}{2}+o(\epsilon)}{s_2(1+t_2\frac{\epsilon}{2s_2}+o(\epsilon))} \\
&= -\frac{(s_1+t_1\frac{\epsilon}{2}+o(\epsilon))(1-t_2\frac{\epsilon}{2s_2}+o(\epsilon))}{s_2} \\
&= -\frac{(s_1+t_1\frac{\epsilon}{2}-s_1t_2\frac{\epsilon}{2s_2}+o(\epsilon))}{s_2} \\
&= -(\frac{s_1}{s_2}+\frac{t_1}{s_2}\frac{\epsilon}{2}-\frac{s_1t_2}{s_2}\frac{\epsilon}{2s_2}+o(\epsilon)) \\
&= -(\frac{s_1}{s_2}+\left(\frac{t_1}{s_2}-\frac{s_1t_2}{s_2^2}\right)\frac{\epsilon}{2}+o(\epsilon))
\end{aligned}$$

**Invasion fitness expansion**

$$\lambda(z, z+\epsilon) = 1 + (\partial\lambda/\partial z)\epsilon + (\partial^2\lambda/\partial z^2)\frac{\epsilon^2}{2} + o(\epsilon^2) \qquad \text{(eq. 5 of maintext)}$$

with $\partial\lambda/\partial z = s_1(z)$ (eq. 6a in maintext) and $\partial^2\lambda/\partial z^2 = (s_2(z) - s_1(z))^2 l^2/(3m\sigma^2) + t_1(z)$ (eq. 6b in maintext).

#### Application to the Laplace case

##### Selection gradient within habitat 1

$$w_1(z) = e^{-\Omega z}$$

$$w_1'(z) = -\Omega e^{-\Omega z}$$

$$w_1''(z) = \Omega^2 e^{-\Omega z}$$

$$s_1(z) = w_1'(z)/w_1(z) = -\Omega$$

$$t_1(z) = w_1''(z)/w_1(z) = \Omega^2$$

##### Selection gradient within habitat 2

$$w_2(z) = e^{-\Omega(1-z)}$$

$$w_2'(z) = \Omega e^{-\Omega(1-z)}$$

$$w_2''(z) = \Omega^2 e^{-\Omega(1-z)}$$

$$s_2(z) = w_2'(z)/w_2(z) = \Omega$$

$$t_2(z) = w_2''(z)/w_2(z) = \Omega^2$$

##### Nagylaki parameters

$$k(z) = \frac{(l/\sqrt{2})\sqrt{\Omega}}{\sigma\sqrt{m}}\sqrt{\epsilon} + o(\sqrt{\epsilon}) \qquad \text{(which leads to eq. 8a of main text)}$$

$$\alpha(z) = 1 - \Omega\epsilon + o(\epsilon) \qquad \text{(which leads to eq. 8b of main text)}$$

**Invasion fitness expansion**

$$\partial\lambda/\partial z = -\Omega$$

$$\partial^2\lambda/\partial z^2 = 4\Omega^2 l^2/(3m\sigma^2) + \Omega^2 = \Omega^2(4l^2/(3m\sigma^2) + 1)$$

$$\lambda = 1 - \Omega\epsilon + \Omega^2(4l^2/(3m\sigma^2) + 1)\frac{\epsilon^2}{2} + o(\epsilon^2)$$

Using $\rho = \frac{\sigma\sqrt{m}}{l\sqrt{\epsilon}}$

$$\begin{aligned}
\lambda &= 1 - \Omega\epsilon + \Omega^2((4/3)l^2/(m\sigma^2) + 1)\tfrac{\epsilon^2}{2} + o(\epsilon^2) \\
&= 1 - \Omega\epsilon + \Omega^2((4/3)1/(\epsilon\rho^2) + 1)\tfrac{\epsilon^2}{2} + o(\epsilon^2) \\
&= 1 - \Omega\epsilon + \Omega^2((4/3)1/\rho^2\epsilon + \epsilon^2)\tfrac{1}{2} + o(\epsilon^2) \\
&= 1 - \Omega\epsilon + \Omega^2((2/3)1/\rho^2\epsilon) + o(\epsilon) \\
&= 1 + ((2/3)\Omega/\rho^2 - 1)\Omega\epsilon + o(\epsilon)
\end{aligned}$$

$$\lambda > 1 \iff (2/3)\Omega/\rho^2 > 1$$

#### Application to the Gauss case

**Selection gradient within habitat 1**

$$w_1(z) = e^{-\Omega^2 z^2}$$

$$w_1'(z) = -2\Omega^2 z e^{-\Omega^2 z^2}$$

$$w_1''(z) = -2\Omega^2(1 - 2\Omega^2 z^2)e^{-\Omega^2 z^2}$$

$$s_1(z) = w_1'(z)/w_1(z) = -2\Omega^2 z$$

$$t_1(z) = w_1''(z)/w_1(z) = -2\Omega^2(1 - 2\Omega^2 z^2)$$

**Selection gradient within habitat 2**

$$w_2(z) = e^{-\Omega^2(1-z)^2}$$

$$w_2'(z) = 2\Omega^2(1-z)e^{-\Omega^2(1-z)^2}$$

$$w_2''(z) = 2\Omega^2(-1 + 2\Omega^2(1-z)^2)e^{-\Omega^2(1-z)^2}$$

$$s_2(z) = w_2'(z)/w_2(z) = 2\Omega^2(1-z)$$

$$t_2(z) = w_2''(z)/w_2(z) = 2\Omega^2(-1 + 2\Omega^2(1-z)^2)$$

#### Nagylaki parameters

$$k(z) = \frac{l\Omega\sqrt{1-z}}{\sigma\sqrt{m}}\sqrt{\epsilon} + o(\sqrt{\epsilon})$$

Therefore $k(0) = \frac{l\Omega}{\sigma\sqrt{m}}\sqrt{\epsilon} + o(\sqrt{\epsilon})$ which leads to eq. 12a of main text.

$$\begin{aligned} \alpha(z) &= -(\tfrac{-z}{1-z} + \left(\tfrac{-(1-2\Omega^2z^2)}{1-z} - \tfrac{z(1-2\Omega^2z^2)}{(1-z)^2}\right)\tfrac{\epsilon}{2} + o(\epsilon)) \\ &= \tfrac{z}{1-z} + \tfrac{(1-2\Omega^2z^2)}{1-z}\left(1 + \tfrac{z}{1-z}\right)\tfrac{\epsilon}{2} + o(\epsilon) \end{aligned}$$

Therefore $\alpha(0) = \epsilon/2 + o(\epsilon)$, which leads to eq. 12b of maintext.

#### Invasion fitness expansion

$$\partial\lambda/\partial z = -2\Omega^2 z$$

Therefore $\partial\lambda/\partial z = 0$ when $z = 0$.

$$\begin{aligned} \partial^2\lambda/\partial z^2 &= (s_2(z) - s_1(z))^2 l^2/(3m\sigma^2) + t_1 \\ &= (2\Omega^2(1-z) + 2\Omega^2 z)^2 l^2/(3m\sigma^2) - 2\Omega^2(1 - 2\Omega^2 z^2) \\ &= (2\Omega^2)^2 l^2/(3m\sigma^2) - 2\Omega^2(1 - 2\Omega^2 z^2) \\ &= 2\Omega^2(2\Omega^2 l^2/(3m\sigma^2) - (1 - 2\Omega^2 z^2)) \end{aligned}$$

Therefore $\partial^2\lambda/\partial z^2 = 2\Omega^2(2\Omega^2 l^2/(3m\sigma^2) - 1)$ when $z = 0$.

$$\begin{aligned} \lambda &= 1 + 2\Omega^2(2\Omega^2 l^2/(3m\sigma^2) - 1)\tfrac{\epsilon^2}{2} + o(\epsilon^2) \\ &= 1 + 2\Omega^2(2\Omega^2 l^2/(3m\sigma^2)\tfrac{\epsilon}{2} - \tfrac{\epsilon}{2})\epsilon + o(\epsilon^2) \\ &= 1 + 2\Omega^2(\Omega^2 l^2/(3m\sigma^2)\epsilon - \tfrac{\epsilon}{2})\epsilon + o(\epsilon^2) \\ &= 1 + 2\Omega^2(k(0)^2/3 - \alpha(0))\epsilon + o(\epsilon^2) \end{aligned}$$

$$\lambda > 1 \iff k(0) > \sqrt{3}\sqrt{\alpha(0)} \qquad \text{(eq. 11 of main text)}$$

### Appendix C : Relative contributions of small and large effect alleles to phenotype after the initial phase of adaptation

#### Establishment probability

Here we assume a situation slightly different form the monomrphic analysis made in Appendix A. We now suppose that the resident genotype is fixed and present two distinct phenotypes $z_1, z_2$ in habitat $1, 2$. We consider a new mutation with phenotypic effect $a$ in both habitats. Denote $\pi(a, z_1, z_2)$ the establishment probability of this mutation. A branching process approximation yields :

$$\pi(a, z_1, z_2) = 2 \max (\Lambda(a, z_1, z_2) - 1, 0)$$

where $\Lambda(a, z_1, z_2)$ is the dominant eigen value of the linearized invasion dynamics. $\Lambda(a, z_1, z_2)$ can be derived from maintext formulas :

$$\Lambda(a, z_1, z_2) = 1 + \Lambda_1(z_1, z_2)a + \Lambda_2(z_1, z_2)\frac{a^2}{2}$$

where $\Lambda_1(z_1, z_2) = s_1(z_1)$, $\Lambda_2(z_1, z_2) = \frac{(s_2(z_2)-s_1(z_1))^2 l^2}{3m\sigma^2} + t_1(z_1)$, $s_i(z) = \frac{w_i^{'}(z)}{w_i(z)}$ for $i \in \{1, 2\}$, and $t_1(z) = \frac{w_1^{''}(z)}{w_1(z)}$. Then :

$$\pi(a, z_1, z_2) = a \max \left( 2s_1(z_1) + a \left( \frac{(s_2(z_2) - s_1(z_1))^2 l^2}{3m\sigma^2} + t_1(z_1) \right), 0 \right)$$

##### Laplace fitness function

With the Laplace fitness function:

$$\begin{aligned}
&z \in (0, 1)\\
&w_1(z) = e^{-\Omega z},\ w_1^{'}(z) = -\Omega e^{-\Omega z}\\
&w_1^{''}(z) = \Omega^2 e^{-\Omega z}\\
&s_1(z) = \tfrac{w_1^{'}(z)}{w_1(z)} = -\Omega\\
&t_1(z) = \tfrac{w_1^{''}(z)}{w_1(z)} = \Omega^2\\
&w_2(z) = e^{-\Omega(1-z)},\ w_2^{'}(z) = \Omega e^{-\Omega(1-z)}\\
&s_2(z) = \tfrac{w_2^{'}(z)}{w_2(z)} = \Omega
\end{aligned}$$

Then $\pi(a, z_1, z_2)$ does not depend on $z_1, z_2$ and:

$$\pi(a, z_1, z_2) = \max \left( a\Omega \left[ a\Omega \left( \frac{4l^2}{3m\sigma^2} + 1 \right) - 2 \right], 0 \right)$$

Given that $a$ is assumed to be small, one needs to assume that $\frac{2\Omega l^2}{3m\sigma^2} \to +\infty$ for polymorphism to emerge. Then:

$$\pi(a, z_1, z_2) \approx 2a\Omega \max\left(a\frac{2\Omega l^2}{3m\sigma^2} - 1, 0\right)$$

##### Gauss fitness function

With Gauss fitness function:

$$\begin{array}{l} z \in (0,1) \\ w_1(z) = e^{-\Omega^2 z^2},\ w_1^{'}(z) = -2\Omega^2 z e^{-\Omega^2 z^2} \\ w_1^{''}(z) = -2\Omega^2(1-2\Omega^2 z^2)e^{-\Omega^2 z^2} \\ s_1(z) = \frac{w_1^{'}(z)}{w_1(z)} = -2\Omega^2 z \\ t_1(z) = \frac{w_1^{''}(z)}{w_1(z)} = -2\Omega^2(1-2\Omega^2 z^2) \\ w_2(z) = e^{-\Omega^2(1-z)^2},\ w_2^{'}(z) = 2\Omega^2(1-z)e^{-\Omega^2(1-z)^2} \\ s_2(z) = \frac{w_2^{'}(z)}{w_2(z)} = 2\Omega^2(1-z) \end{array}$$

Then :

$$\pi(a, z_1, z_2) = 2a\Omega^2 \max\left(2a\Omega^2\left(\frac{(1-z_2+z_1)^2 l^2}{3m\sigma^2} + z_1^2\right) - a - 2z_1, 0\right)$$

Given that $a$ is assumed to be small, one needs to assume that $z_1$ is also small for polymorphism to emerge. Then:

$$\pi(a, z_1, z_2) \approx 2a\Omega^2 \max\left(a\left(\frac{2\Omega^2 l^2}{3m\sigma^2}(1-z_2)^2 - 1\right) - 2z_1, 0\right)$$

#### Approximating trait dynamics

##### Assumptions

Mutants with a small phenotypic effect $a$ occurs in the system at a rate $nN\mu_a$ where $\mu_a$ is the *per capita* mutation rate, $n$ the number of demes and $N$ the number of individuals in each deme. Similarly, mutants with a large phenotypic effect $A$ occur in the system at a rate $nN\mu_A$.

Therefore, when there is a resident monomorphic genotype with phenotype $z_i$ in habitat $i$, $i \in \{1, 2\}$, the establishment rates of new small and large effect alleles are $nN\mu_a\pi(a, z_1, z_2)$ and $nN\mu_A\pi(A, z_1, z_2)$ respectively.

When a mutant invades, it should create a cline in the landscape. Here we simplify this phenomenon by assuming that the mutant invasion creates a shift in the trait of the monomorphic population of $b$ in habitat 2 and $\rho b, \rho < 1$ in habitat 1, where $b \in \{a, A\}$ is the effect size of the mutation.

The initial state of the system is $(z_{in}, z_{out}) = (0, 0)$, therefore with our simplifying assumptions we expect $z_{out} = \rho z_{in}$ at all times and we focus on $z_{in}$ dynamics and we note $z = z_{in}$.

##### Trait dynamics within pocket

With the above assumptions and using a continuous-time approximation, the expected value of $z$ should follow the dynamics:

$$z' = 2nN\left[\mu_a \pi(a, z_1, z_2)a + \mu_A \pi(A, z_1, z_2)A\right]$$

Denote $z_a(t)$ (resp. $z_A(t)$) the contribution of small (resp. large) alleles to the trait at time $t$. Then $z(t) = z_a(t) + z_A(t)$ and:

$$\begin{aligned} z_a' &= 2nN\mu_a \pi(a, z_1, z_2)a \\ z_A' &= 2nN\mu_A \pi(A, z_1, z_2)A \end{aligned}$$

#### Deriving small alleles relative contribution to the trait

Denote $Q_a(t) = \frac{z_a}{z}$ which approximately corresponds to the expected proportion of the trait generated by small alleles. The dynamics of $Q_a(t)$ is defined by the initial condition :

$$Q_a(0) = \frac{\mu_a \pi(a, 0, 0)a}{\mu_a \pi(a, 0, 0)a + \mu_A \pi(A, 0, 0)A}$$

and the ordinary differential equation :

$$Q_a'(t) = \frac{z_a'}{z} - \frac{z'}{z} Q_a$$

Noting that $z(t)$ is monotically increasing with time, we can define $\tilde{Q}_a(z)$ such that $\tilde{Q}_a(z(t)) = Q_a(t)$. Then, noting that $\tilde{Q}_a'(z) = \frac{Q_a'}{z'}$, the dynamics of $\tilde{Q}_a$ follows the ordinary differential equation :

$$\tilde{Q}_a'(z) + \frac{1}{z}\tilde{Q}_a = \frac{1}{z}\frac{\mu_a \pi(a, \rho z, z)a}{\mu_a \pi(a, \rho z, z)a + \mu_A \pi(A, \rho z, z)A}$$

The ODE has the following solution :

$$\tilde{Q}_a(z) = \frac{1}{z}\int_0^z \left(\frac{\mu_a \pi(a, \rho u, u)a}{\mu_a \pi(a, \rho u, u)a + \mu_A \pi(A, \rho u, u)A}\right) du = \frac{1}{z}\int_0^z \left(\frac{1}{1 + \frac{\mu_A}{\mu_a}\frac{A}{a}\frac{\pi(A,\rho u,u)}{\pi(a,\rho u,u)}}\right) du$$

##### Laplace fitness function

If $A < \frac{3m\sigma^2}{2\Omega l^2}$ local adaptation cannot occur. Assuming $A > \frac{3m\sigma^2}{2\Omega l^2}$, $Q_a$ is constant through time and one expects at the end of the initial bout of adaptation :

$$\lim_{t \to +\infty} Q_a(t) = \begin{cases} 0 & \text{if } a < \frac{3m\sigma^2}{2\Omega l^2} < A \\ \frac{1}{1 + \frac{\mu_A}{\mu_a}\left(\frac{A}{a}\right)^2 \frac{\left[A - \frac{3m\sigma^2}{2\Omega l^2}\right]}{\left[a - \frac{3m\sigma^2}{2\Omega l^2}\right]}} & \text{if } \frac{3m\sigma^2}{2\Omega l^2} < a \end{cases}$$

Above results means that when $a < \frac{3m\sigma^2}{2\Omega l^2} < A$, only large alleles can invade and the contribution of small alleles to the phenotype within the pocket remain null. When $l$ increases in such a way that $\frac{3m\sigma^2}{2\Omega l^2} < a$, small alleles can invade and their relative contribution to adaptive phenotype increases with $l$. When $l \to +\infty$, the relative contribution tends towards $\frac{1}{1+\frac{\mu_A}{\mu_a}\left(\frac{A}{a}\right)^3}$.

**Gauss fitness function**

If $1 < \frac{3m\sigma^2}{2\Omega^2 l^2}$ local adaptation cannot occur. Assuming $1 > \frac{3m\sigma^2}{2\Omega^2 l^2}$,

$$\tilde{Q}_a(z) = \begin{cases} \frac{1}{z}\int_0^z \left( \frac{1}{1+\left[\frac{\mu_A}{\mu_a}\left(\frac{A}{a}\right)^3\right] \frac{\frac{2\Omega^2 l^2}{3m\sigma^2}(1-u)^2-2\frac{\rho}{A}u-1}{\frac{2\Omega^2 l^2}{3m\sigma^2}(1-u)^2-2\frac{\rho}{a}u-1}} \right) du & \text{if } z < u_0 \\ \frac{1}{z}\int_0^{u0} \left( \frac{1}{1+\left[\frac{\mu_A}{\mu_a}\left(\frac{A}{a}\right)^3\right] \frac{\frac{2\Omega^2 l^2}{3m\sigma^2}(1-u)^2-2\frac{\rho}{A}u-1}{\frac{2\Omega^2 l^2}{3m\sigma^2}(1-u)^2-2\frac{\rho}{a}u-1}} \right) du & \text{if } z > u_0 \end{cases}$$

where $u_0$ is the unique root of $\frac{2\Omega^2 l^2}{3m\sigma^2}(1-u_0)^2 - 2\frac{\rho}{a}u_0 - 1$ on $(0,1)$, well defined when $1 > \frac{3m\sigma^2}{2\Omega^2 l^2}$. From this equation we deduce that $\tilde{Q}_a$ is not necessarily monotonous during the adaptation process but always remains below its initial value. In particular, when $z$ becomes greater than $u_0$ the relative contribution of small alleles decreases since only large effect alleles can further contribute to adaptation. At the end of this bout of adaptation, one expects the phenotypic contribution of small alleles to be :

$$\lim_{t\to+\infty} Q_a(t) = \begin{cases} \text{undefined} & \text{if } 1 < \frac{3m\sigma^2}{2\Omega^2 l^2} \\ \frac{1}{u_1}\int_0^{u0} \left( \frac{1}{1+\frac{\mu_A}{\mu_a}\left(\frac{A}{a}\right)^3 \frac{\frac{2\Omega^2 l^2}{3m\sigma^2}(1-u)^2-2\frac{\rho}{A}u-1}{\frac{2\Omega^2 l^2}{3m\sigma^2}(1-u)^2-2\frac{\rho}{a}u-1}} \right) du & \text{if } \frac{3m\sigma^2}{2\Omega^2 l^2} < 1 \end{cases}$$

where $u_1 > u_0$ is the unique root of $\frac{2\Omega^2 l^2}{3m\sigma^2}(1-u_1)^2 - 2\frac{\rho}{A}u_1 - 1$, which is well defined when $1 > \frac{3m\sigma^2}{2\Omega^2 l^2}$

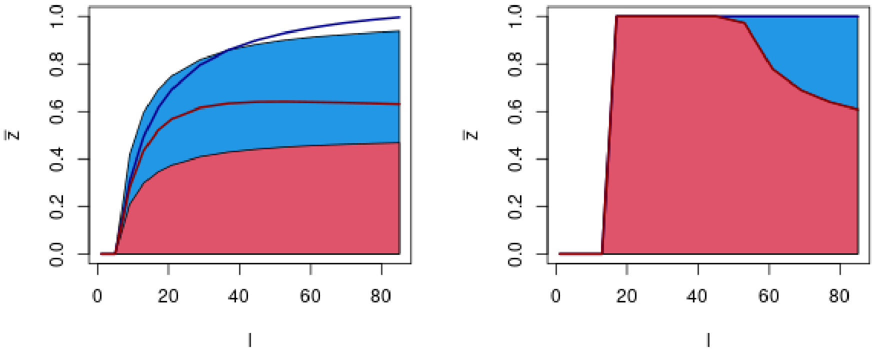


Figure 1: Theoretical prediction of the relative contribution of small and large effect alleles to the phenotype at the center of habitat 2 after the initial phase of adaptation. We used main text parameters : $m = 0.5, d = 10, \Omega = 1, \mu_a = 10^{-5}, \mu_A = 10^{-8}, a = 0.01, A = 0.1$. Gauss and Laplace fitness functions are presented on the left and right panels respectively. Shapes correspond to $\rho = 0$ (i.e. mutations only change the phenotype in environment 2; see section **Assumptions**). Blue and red shapes present how small and large alleles respectively contribute to the total phenotype at the middle of habitat 2 for various sizes of habitat 2 ($l$). Dark blue and red lines show how the frontiers of these shapes are modified when $\rho = 0.1$ (i.e. mutations also change the phenotype in environment 1, but the change in only 10% of the one observed in environment 2; see section **Assumptions**). In particular, the contribution of large alleles becomes non-monotonic with $l$ with Gauss fitness function.
